# Shared binding sites for the chromosomal architectural protein Su(Hw) mediate physical interactions between *Drosophila* TAD boundaries

**DOI:** 10.64898/2026.05.28.727987

**Authors:** Wenfan Ke, Miki Fujioka, Brendan Wang, Tiffany Park, Li Zhang, Amina Kurbidaeva, Yuri Pritykin, James B Jaynes, Paul Schedl

**Author notes:** Laboratory of Dairy Science of the Education Ministry, Northeast Agricultural University, Harbin 150030, Heilongjiang, P.R China. Center for Systems Biology, New York University, New York, NY 10003.

## Abstract

The head-to-tail pairing of the *nhomie* and *homie* boundaries is responsible for the formation of a stem-loop TAD (topologically associated domain) that encompasses the *even-skipped* (*eve*) gene and its regulatory elements. TAD boundaries like *homie* and *nhomie* have partner preferences and one mechanism thought to be important in linking compatible boundaries is shared binding sites for the same architectural protein. Both *eve* boundaries have a single binding site for the polydactyl zinc finger protein Su(Hw). Here we have investigated the role of the shared Su(Hw) sites in the long-distance regulation of a dual reporter by the *eve* enhancers. Consistent with this mechanism, we find that when the transgene *nhomie* or *homie* boundary has a mutation in the Su(Hw) binding site, reporter activation by the *eve* enhancers is severely compromised. Using MicroC we show the mutating the single Su(Hw) substantially weakens the ability of the *nhomie* and *homie* boundaries in the transgene to physically pair with the two endogenous *eve* boundaries. Su(Hw) binding sites are not, however, sufficient as the *gypsy* su(Hw) insulator is unable to support long-distance regulation. In a less demanding transvection assay mutations in the *nhomie* and *homie* Su(hw) site have only a modest effect on regulatory interactions. The *eve* boundaries can mediate transvection with the *gypsy* insulator; however. they are not compatible with the dCTCF *Fab-8* boundary even in the transvection assay. Finally, we show that the regulatory and physical interactions of the *nhomie* boundary in the transgene and the two endogenous *eve* boundaries are incompatible with the popular cohesin-loop extrusion model.

## Introduction

Studies on lampbrush chromosomes were the first to suggest that the key organizing principle for chromosomes in animals is the subdivision of the chromatin into a series of topologically independent loops (now called TADs) that extend from the main chromosomal axis (Gall and Callan, 1962; Callan, 1986; Callan, 1987). Each lampbrush chromosome has a stereotypic pattern of loops, and in most loops there is only a single actively transcribed gene. The loops are anchored to the main axis of the chromosome, and the anchors pull apart when the chromosomes are stretched (Callan, 1986, 1987). Since then, a variety of approaches have shown that TADs are pervasive features of chromosomes in multicellular organisms (Rao et al., 2014; Dekker and Heard, 2015; Eagen et al., 2015; Stadler et al., 2017; Hsieh et al., 2021; Krietenstein et al., 2020; Goel et al., 2023; Dolstein et al., 2025). In flies, TADs range in size from only a few kb to over 100 kb, while recent MicroC and Region Capture MicroC (RCMC) studies indicate that the scale is similar in mammals. The arrangement of TADs along a given chromosome tends to be invariant and is largely, but not completely, independent of cell type or developmental stage (Bing et al. 2024; Ke et al., 2024; Dolstein et al., 2025; Wang et al., 2026).

This regular and heritable organization is generated by a special class of *cis-*acting elements called boundaries or insulators. Boundary elements were first discovered and have been most fully characterized in *Drosophila*; however, similar elements have been identified in other species (Ghirlando et al., 2012; Ghirlando and Felsenfeld, 2016; Chetverina et al., 2017; Matthews and White 2019; Cavaleiro et al., 2021). Boundary elements in flies consist of one or more large (>150 bp) nuclease hypersensitive regions and can span DNA sequences of up to 1.5 kb in length. These nuclease-hypersensitive regions are targets for a large and diverse collection of DNA binding proteins that have been implicated in boundary function (Parkhurst et al., 1988; Gasner et al., 1999; Golovin et al., 2007; Maksimento et al., 2015; Zolotarev et al, 2016; Fedotova et al., 2017; Chetverina et al. 2021; Bonchuk et al., 2021; Bonchuk et al., 2023). Of these, the most prevalent class are polydactyl C2H2 zinc finger proteins {e.g., CTCF, Zw5, Su(Hw), Pita, Zipic, Clamp, Zad1}(Bonchuk et al., 2021; Fedotova et al., 2017). Other boundary factors include several BEN domain proteins (Insv, Elba1-2), GAF, Ibf1/2, BEAF, CP190 and the 31 Mod(mdg4) isoforms (Zhao et al., 1995; Bonchuk et al., 2011; Aoki et al., 2012; Dai et al. 2013; Cuartero et al., 2013; Avva and Hart, 2016). As there are over 200 mostly uncharacterized polydactyl C2H2 zinc finger proteins, including some 70 proteins that have the ZAD homodimerization domain, encoded in the fly genome (Bonchuk et al., 2021), it is likely that a large number of chromosomal architectural proteins have yet to be identified.

In flies, TADs are formed by boundary:boundary pairing. The first evidence of physical pairing came from experiments showing that when either the Bithorax (BX-C) boundary *Mcp* or the *gypsy* transposon *su(Hw)* boundary are included in transgenes, they can induce long-distance regulatory interactions (PcG-dependent silencing and enhancer activation) between transgene inserts located many megabases (Mb) apart (Vazquez et al., 1993; Sigrest and Pirrotta, 1997; Muller et al., 1998). Regulatory interactions were even observed for transgenes inserted on different chromosomes. Subsequent *in vivo* imaging studies showed that 2, 3, and even 4 copies of these distant transgenes co-localize in imaginal disc cells (Vazquez et al., 2006; Li et al., 2011). The parameters governing boundary pairing interactions have been defined using boundary bypass, transvection and boundary competition assays (Cai and Shen, 2001; Muravyova et al., 2001; Gohl et al., 2011; Li et al., 2018). Though boundary elements can engage in promiscuous pairing interactions, there are clear partner preferences. For example, in boundary bypass assays using boundaries from the *Abd-B* region of BX-C, Kyrchanova et al. (2011) found that *Mcp* can pair with *Fab-8* but not with *Fab-7*. The other key feature is orientation dependence (Kyrchanova et al., 2008a). With a few exceptions, pairing interactions are orientation-dependent. Orientation dependence can differ depending upon whether boundaries are pairing with themselves or with other boundaries. The self-pairing interactions that have been examined in detail are head-to-head (Kyrchanova et al., 2008a; Kyrchanova et al., 2008b; Fujioka et al., 2016). Self-pairing interactions take place in *trans*, and these interactions are likely responsible for the alignment and pairing of sister chromosomes and homologs (Fujioka et al., 2016; Alhai Abed et al., 2019; Viets et al., 2019; Child et al., 2021). That these interactions are head-to-head is not surprising, as head-to-tail self-pairing would generate unpaired loops and potentially disrupt transvection (Fujioka et al., 2016). Heterologous pairing interactions typically take place in *cis*, and unlike self-pairing, can be either head-to-head or head-to-tail. The former generates a TAD with a circle-loop topology while the latter generates a TAD with a stem-loop topology (Kyrchanova et al., 2008a; Fujioka et al., 2016; Li et al., 2018; Bing et al., 2024; Ke et al., 2024).

Thus far, three different mechanisms are thought to be involved in boundary:boundary pairing interactions. The simplest mechanism in flies is based on shared binding sites for architectural proteins that can form dimers or multimers (Bonchuk et al. 2021; Fedotova et al. 2017). Most of the factors implicated in fly boundary function form such dimers or higher order multimers. A second mechanism for physically linking two boundaries is protein:protein interaction between heterologous DNA binding factors (Blanton et al. 2003). Yet a third mechanism would be proteins that function to bridge DNA binding proteins that are associated with each boundary. Several proteins that can potentially function as “linkers” have been identified. Included in this group are two proteins, CP190 and the 67.1 isoform of Mod(mdg4), which interact with the polydactyl zinc finger protein Su(Hw) (Harrison et al.,1993; Kim et al., 1996; Pai et al., 2004; Golovnin et al., 2007; Vogelmann et al., 2014; Melnikova et al., 2017; Melnikova et al., 2018; Melnikova et al., 2019; Kaushal et al., 2022; Golovnin et al., 2023). CP190 and Mod(mdg4) family proteins, including the 67.1 isoform, have BTB domains that assemble into dimers (CP190) or multimers (Mod(mdg4)), and thus could form a bridge linking Su(Hw) proteins bound to sites in different boundaries.

In the studies reported here, we have tested the role of the chromosomal architectural protein Su(Hw) in generating direct physical interactions between boundary elements. As a model system for this analysis we used the two boundaries, *nhomie* and *homie*, that are responsible for the formation of the ∼16 kb TAD that encompasses the *Drosophila even-skipped* (*eve*) gene (Fujioka et al., 2009; Fujioka et al., 2016; Ke et al., 2024). Previous work showed that *nhomie* and *homie* pair with themselves head-to-head, while they pair with each other head-to-tail (Fujioka et al., 2016). Like the *Mcp* and *su(Hw)* boundaries, *nhomie* and *homie* are also able to mediate long-distance regulatory interactions. We have taken advantage of this long-distance activity to probe the pairing interactions of the two *eve* boundaries. In one experimental paradigm, we used an attP site located about a dozen TADs upstream from *eve* (at -142 kb, in the 1^st^ exon of the *hebe* gene) to introduce a transgene containing two reporters driven by the *eve* promoters, *eve-LacZ* and *eve-GFP* (see Figure 1). When *nhomie* or *homie* is inserted between the two reporters, one of the two reporters is activated more strongly by *eve* enhancers than the other. Activation depends on the 5’→3’ orientation of the boundary relative to the two reporters in the transgene, but not on the orientation of the transgene in the chromosome. Thus, *eve-LacZ* is preferentially activated when it is positioned “downstream” of *nhomie* or “upstream” of *homie*. The same relationship holds for *eve-GFP*. We also found that the ability of a minimal 271 bp *homie* fragment (DEF) to mediate *eve* enhancer-dependent reporter activation in the -142 kb assay depends on a predicted Su(Hw) binding site located in the D element (Fujioka et al., 2025). Since *nhomie* also has a predicted Su(Hw) binding site, this shared DNA recognition sequence could serve two functions: namely *nhomie* and *homie* self-pairing and heterologous pairing. In the studies reported here we have tested this idea. We have also analyzed interactions between the *gypsy* transposon *Su*(*hw*) insulator and the two *eve* boundaries.

**Figure 1.**
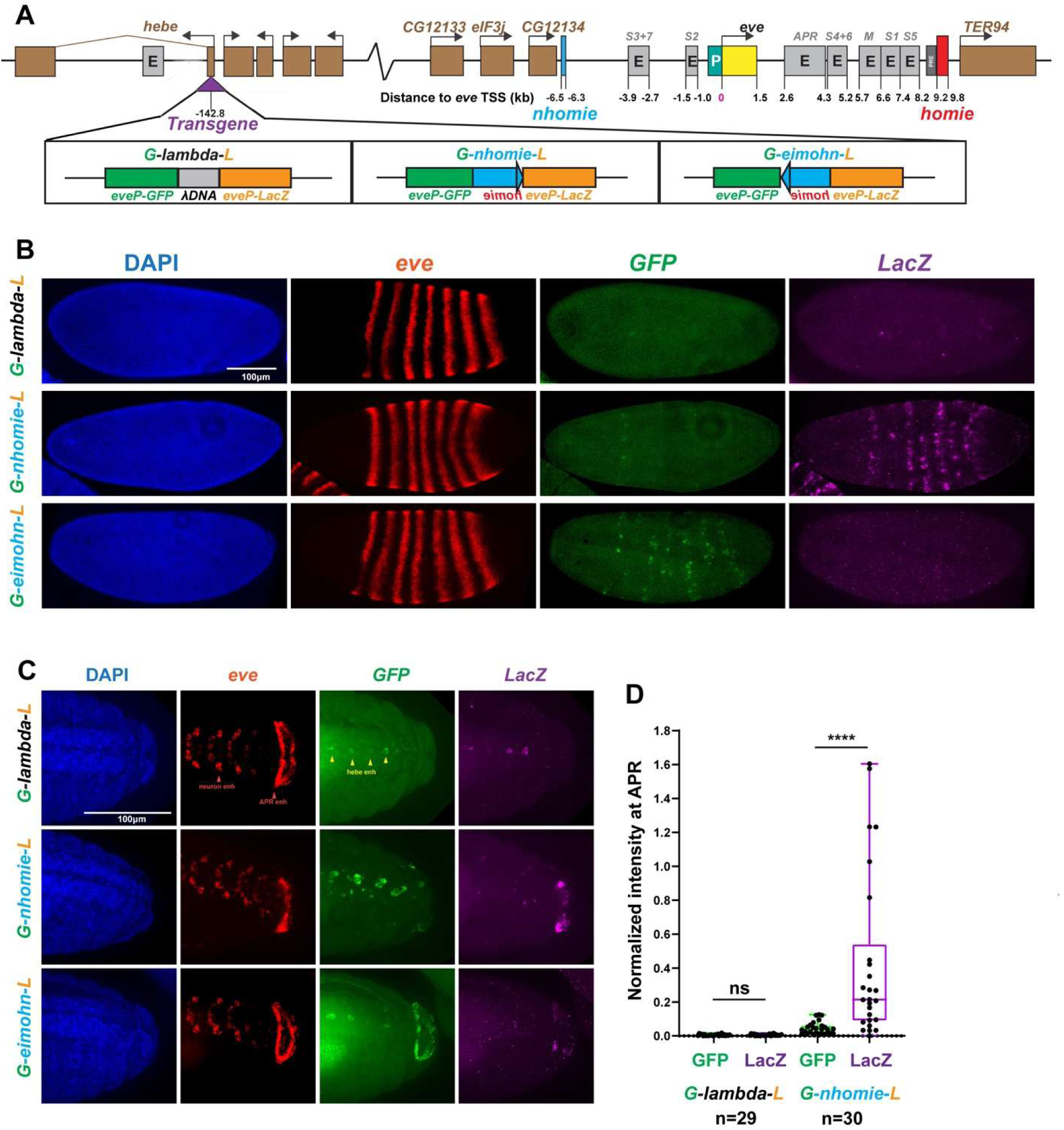
Activation by *eve* enhancers of *nhomie*-containing transgenes inserted at -142 kb from *eve*. A. Map of *eve* locus, a subset of the intervening genes and the attP site in the first exon of the *hebe* gene. The three different transgene inserts oriented so that the *LacZ* reporter is closest to *eve* are shown: *lambda* DNA (*G-lambda-L*), *nhomie* in the same (forward) orientation as the *eve nhomie* (*G-nhomie-L*), and *nhomie* in the opposite (reverse) orientation as the *eve nhomie* (*G-eimohn-L*). B) The panels in each row show DAPI staining, *eve* mRNA, *GFP* mRNA, and *LacZ* mRNA for *G-lambda-L*, *G-nhomie-L* and *G-eimohn-L* blastoderm stage embryos. C) The panels in each row show DAPI staining, *eve* mRNA, *GFP* mRNA, and *LacZ* mRNA for *G-lambda-L*, *G-nhomie-L* and *G-eimohn-L* in stage 13-14 embryos. D) Quantitation of transgene expression (*LacZ* or *gfp*) for *G-lambda-L*, *G-nhomie-L*, and *G-eimohn-L* in stage 13-14 embryos.

## Results

### nhomie-dependent reporter activation

To assay the long-distance pairing activity of the *nhomie* boundary, we inserted a “minimal” (602 bp) version of *nhomie* into the dual reporter in each orientation. The resulting transgenes were then integrated into the -142 kb *attP* site to give four different transgene configurations: *G-nhomie-L* and *G-eimohn-L* (Figure 1 and Figure 1-figure supplemental 1), and *L-nhomie-G* and *L-eimohn-G* (Figure 1-figure supplemental 2). In the transgenes *G-nhomie-L* and *G-eimohn-L* (diagrammed in Figure 1A), the *LacZ* reporter is located between the transgene *nhomie* and the *eve* TAD. Conversely, in the transgenes *L-nhomie-G* and *L-eimohn-G* (shown in Figure 1-figure supplemental 2), the *GFP* reporter is located between the transgene *nhomie* and the *eve* TAD. Upstream of the *attP* insertion site are enhancers for the *hebe* gene, which are active in stage 12-16 embryos (Fujioka et a., 2025). Shown in Figure 1 is a negative control transgene containing phage *lambda* DNA sequences (*G-lambda-L*) instead of *nhomie*. It is oriented so that *LacZ* is on the *eve* side of the *lambda* DNA, while *GFP* is on the same side as the *hebe* enhancers.

In the transgene *G-nhomie-L* (shown in Figure 1B), *nhomie* is oriented so that the *LacZ* reporter is “downstream” of the boundary, just like the *eve* gene is “downstream” of the endogenous *nhomie* boundary in the *eve* TAD. In this configuration, *LacZ* (but not *GFP*) is preferentially activated by the *eve* enhancers (Fujioka et al., 2016). At the blastoderm stage, the *eve* stripe enhancers drive *LacZ* expression (Fig 1B and Figure 1-figure supplemental 1A), while in stage 13 embryos, *LacZ* expression is driven by the *eve* neurogenic, mesodermal, and anal plate ring (APR) enhancers (Figure 1C and Figure 1-figure supplemental 1A). Quantification of *LacZ* and *GFP* expression in the APR confirmed the preferential expression of *LacZ* (Figure 1D, *G-nhomie-L*). Activation of the *LacZ* reporter by the *eve* enhancers depends upon the presence of *nhomie*, as the *LacZ* reporter is not activated by the *eve* enhancers in the *lambda* DNA control at either stage (Figure 1B and C). Reporter activation also depends upon the orientation of *nhomie* in the transgene. As shown in Figure 1B and C (also see Figure 1-figure supplemental 1A), the *eve* enhancers drive very little *GFP* expression from the *G-nhomie-L* transgene. However, since the *GFP* reporter is on the same side of *nhomie* as the *hebe* enhancers, it is activated by the *hebe* enhancers in stage 13 embryos. The *GFP* reporter is also activated by the *hebe* enhancers in the *lambda* DNA control (Figure 1A); however, the control differs from *G-nhomie-L* in that the *LacZ* reporter is not insulated from the *hebe* enhancer, and they also activate its expression.

Reporter activation depends upon the orientation of the *nhomie* boundary with respect to the reporters in the transgene, and not on the orientation of the transgene in the chromosome. In *G-eimohn-L*, *LacZ* is still on the *eve* side of *nhomie* while *GFP* is located on the other side. However, since *nhomie* is inverted, the *GFP* reporter is now “downstream” of the transgene boundary. In this configuration the *GFP* reporter is activated by the *eve* stripe enhancers at the blastoderm stage (Figure 1B and Figure 1-figure supplemental 1B), and by the tissue-specific enhancers in stage 13 embryos (Figure 1C and Figure 1-figure supplemental 1B). It will also be activated by the *hebe* enhancers, while the *LacZ* reporter is not. Quantification of *LacZ* and *GFP* expression in the APR confirmed this preferential expression of *GFP* rather than *LacZ* (Figure 1D, *G-eimohn-L*).

An analogous set of results is observed for the two transgenes *L-nhomie-G* and *L-eimohn-G*, in which *GFP* is on the *eve* side of the *nhomie*, and *LacZ* is on the *hebe* enhancer side. For *L-nhomie-G*, the *eve* enhancers activate *GFP* while *LacZ* is regulated by the *hebe* enhancer, since *GFP* is in the “downstream” position (Figure 1-figure supplemental 2A). When the orientation of *nhomie* is flipped to give *L-eimohn-G* (*LacZ* is now in the “downstream” position), *LacZ* is preferentially activated by both the *eve* and *hebe* enhancers, while *GFP* is not (Figure 1-figure supplemental 2B).

While orientation-dependent pairing of the transgene *nhomie* with the endogenous *nhomie* and *homie* boundaries results in the preferential activation of the “downstream” reporter, the close physical proximity of the reporter on the “upstream” side of the transgene *nhomie* to the enhancers in the *eve* TAD can also trigger its activation. Activation of the upstream reporter is quite weak (see Figure 1) and is best visualized when *LacZ* is “upstream” of the transgene *nhomie*, and the digoxigenin procedure is used for *in situ* hybridization. This is shown for the *G-nhomie-L* and *G-eimohn-L* pair in Figure 1-figure supplemental 1. As was observed with HCR-FISH probes, only *LacZ* transcripts are detected in *G-nhomie-L* embryos using digoxigenin *in situ* hybridization. However, when *GFP* is downstream of the transgene *nhomie* in *G-eimohn-L* and should be activated by the *eve* enhancers, we detect not only *GFP* but also *LacZ* transcripts. The *LacZ* probe in the digoxigenin experiment is ∼2.5x longer than the *GFP* probe, and this difference likely explains why *LacZ* transcripts are observed in *G-eimohn-L* embryos. Similar results are obtained for the second pair of *nhomie* inserts (*L-nhomie-G* and *L-eimohn-G*): when *GFP* is downstream of the transgene *nhomie*, a low level of *LacZ* transcripts is detected (Figure 1-figure supplemental 2).

### Physical interactions between the reporters at -142 kb and the eve TAD

We used MicroC to probe the physical interactions between the *nhomie* transgenes at -142 kb and sequences in the *eve* TAD. Figure 2A shows the MicroC contact profile in the chromosomal segment between the *eve* TAD and the *attP* site in the 1^st^ exon of the *hebe* gene, as well as a blowup of both the *eve* TAD and the insertion site. Note that there are multiple TADs in the 142 kb chromosomal segment between *eve* and the insertion site at the beginning of the *hebe* gene. As reported previously for the *lambda* DNA control (Bing et al. 2024), weak physical interactions can be detected in 12-16 hr embryos collected at 25°C (Figure 2B and 2C) even though we do not observe activation of either reporter by the *eve* enhancers (Figure 1A). We suspect that the weak contacts seen for the *lambda* control may be mediated by the two *eve* promoters in the transgene.

**Figure 2.**
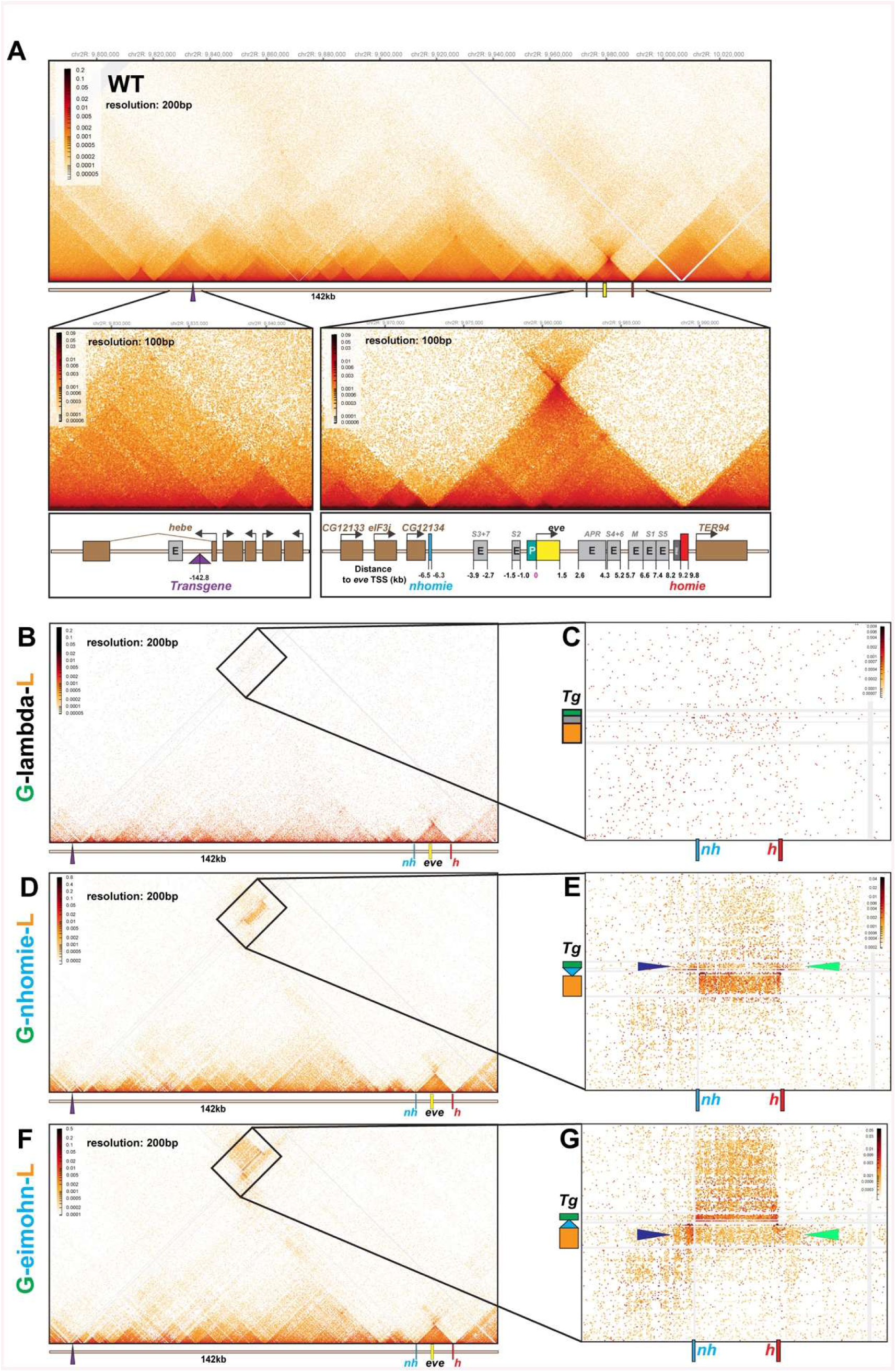
*nhomie*-containing transgenes physically interact with the *eve* TAD. A) Top panel shows the MicroC contact profile in the wild-type chromosomal segment that contains both the attP transgene insertion site (in the *hebe* gene) and the *eve* TAD. Bottom panels show blowups of the MicroC contact profile for the vicinity of the attP site in the *hebe* gene and the *eve* TAD. Note that the MicroC contact profile for the *eve* TAD has the characteristic signature of a stem-loop TAD—a plume above the volcano triangle. B) and C) MicroC contact profile and blow up for the *G-lambda-L* transgene. D) and E) MicroC contact profile and blowup for the *G-nhomie-L* transgene. Blue and green arrowheads indicate interactions between the *GFP* reporter and the TADs flanking the *eve* TAD. F) and G) MicroC contact profile and blow up for the *G-eimohn-L* transgene. Blue and green arrowheads indicate interactions between the *LacZ* reporter and the TADs flanking the *eve* TAD.

A different result is observed for the two *nhomie-*containing transgenes, *G-nhomie-L* and *G-eimohn-L*. In both cases, the physical interactions in 14-16 hr embryos recapitulate the pattern of activity of the transgene reporters. For *G-nhomie-L,* the *LacZ* reporter preferentially interacts with sequences in the *eve* TAD. This interaction between the transgene and *eve* is shown in Figure 2D and E. As can be seen in the blow-up, the frequency of crosslinking events between *LacZ* and sequences in the *eve* TAD is much higher than those between *GFP* and the *eve* TAD.

While the *LacZ* reporter preferentially interacts with sequences in the *eve* TAD, contacts between the *GFP* reporter and the *eve* TAD are much more frequent than those observed for either reporter in the *lambda* control (compare Figure 2B and C with Figure 2D and E). This difference is likely due to the fact that *nhomie* interactions with *eve nhomie* and *homie* brings the *GFP* reporter into sufficiently close proximity for it to interact with sequences in the *eve* TAD, even though the pairing orientation of *nhomie* with the *eve* boundaries favors interactions with *LacZ*.

When *nhomie* is inverted in the transgene to give *G-eimohn-L*, the pattern of physical interactions between the reporters and sequences in the *eve* TAD is the opposite of that observed for *G-nhomie-L*. The primary interactions are between the *GFP* reporter and the *eve* TAD while the secondary interactions are between *LacZ* and the *eve* TAD (Figure 2F and G). This can be seen by comparing the blowups for Figure 2E and 2G.

The distinct topology of the loops generated when the *nhomie* elements in *G-nhomie-L* and *G-eimohn-L* interact with *nhomie* and *homie* in the *eve* TAD results in other differences in the contact patterns. For example, in the *G-nhomie-L* insert, *GFP* comes into contact with the TADs immediately upstream (blue arrowhead in Figure 2E) and downstream (green arrowhead) of the *eve* TAD. When the orientation of *nhomie* is flipped so that the *GFP* reporter is activated instead of *LacZ*, the interaction pattern with neighboring sequences changes. In this case, it is *LacZ* that interacts with sequences in the TADs upstream (blue arrowhead in Figure 2G) and downstream (green arrowhead) of the *eve* TAD.

### Viewpoints from the LacZ and GFP reporters

To further document that boundary orientation in the transgene determines which of the two reporters preferentially interacts with the *eve* TAD, we used the MicroC data to generate “virtual 4C” viewpoints from the *GFP* reporter (Figure 3A) or *LacZ* reporter (Figure 3B) in the *G-nhomie-L* insert, and the *GFP* (Figure 3C) and *LacZ* (Figure 3D) reporters for the *G-eimohn-L* insert. These “virtual 4C” data can also be visualized as 1-dimensional read coverage tracks and the average contact counts between the reporters and the *eve* locus are shown in Figure 3E. For the *G-nhomie-L* insert, the *LacZ* reporter interacts strongly with sequences within the *eve* TAD, while there is little interaction with sequences in TADs located beyond either the *nhomie* or *homie* boundaries. While the *GFP* reporter also interacts with sequences in the *eve* TAD, the interactions are highest near the boundaries, and lower within the *eve* TAD. Additionally, the *GFP* reporter contacts sequences in the TADs flanking the *eve* gene. A similar pattern is observed for *G-eimohn-L*, except in this case, *GFP* preferentially interacts with the *eve* TAD, and the *LacZ* reporter contacts areas flanking the *eve* TAD. The different topologies of the loops generated by interaction between *nhomie* in *G-nhomie-L* and *G-eimohn-L* and *nhomie* / *homie* in the *eve* TAD are shown in Figure 3F.

**Figure 3.**
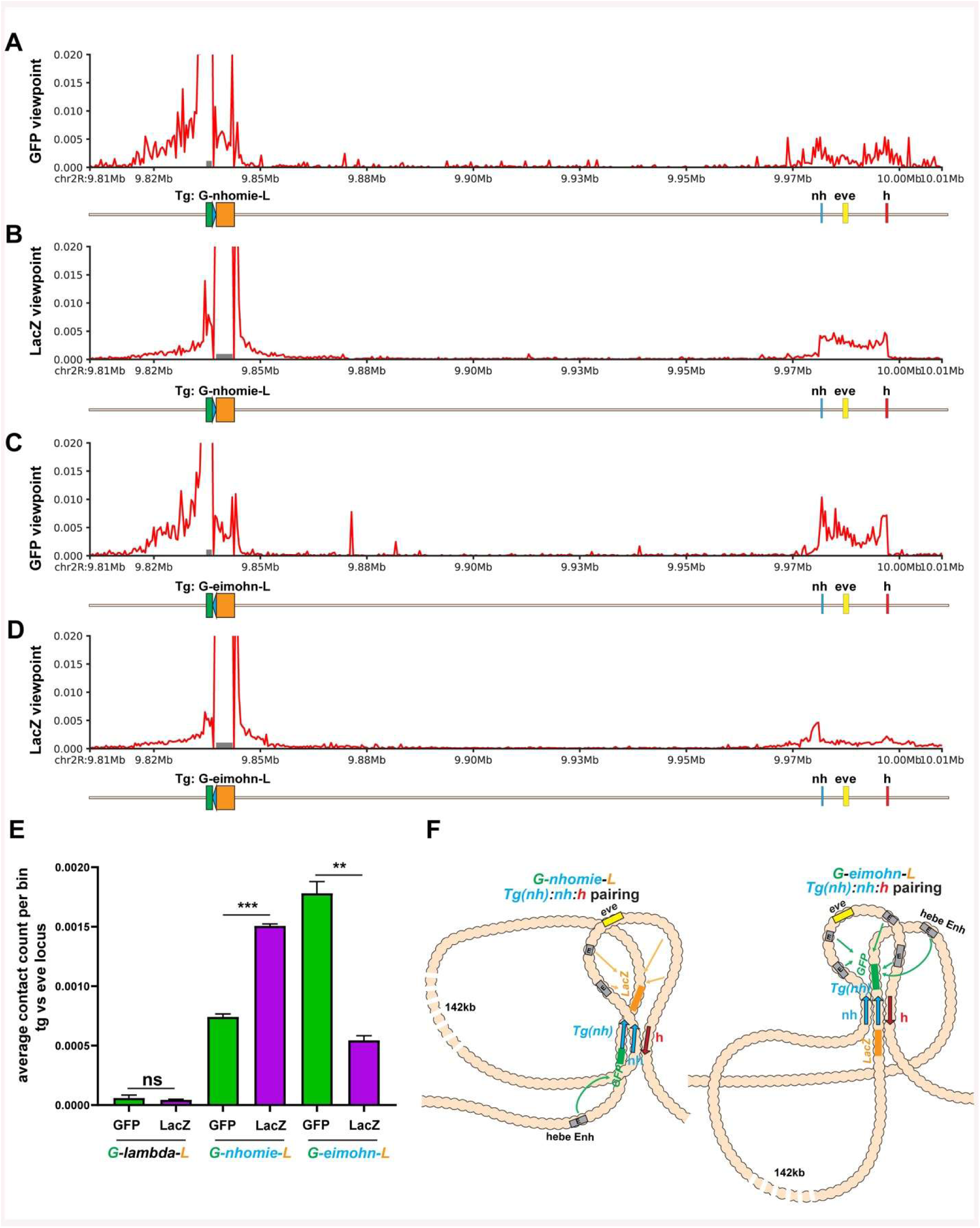
MicroC viewpoints from the *G-nhomie-L* and *G-eimohn-L* reporters. Viewpoints for the *G-nhomie-L* transgene for A) *GFP* and B) *LacZ* reporters. Viewpoints for the *G-eimohn-L* transgene for C) *GFP* and D) *LacZ* reporters. E) The average contact count per bin for the *GFP* and *LacZ* reporters, as indicated for the three transgene inserts *G-lambda-L*, *G-nhomie-L*, and *G-eimohn-L*. F) Models for chromatin organization of *G-nhomie-L* and *G-eimohn-L*.

### The Su(Hw) site in nhomie is required for long-distance regulatory interactions

Fujioka et al. (2025) showed that the predicted Su(Hw) binding site in a minimal 271 bp *homie* boundary (called DEF) is required for mediating long-distance regulatory interactions in the -142 kb assay. Since *nhomie* also has a predicted Su(Hw) binding site, we tested whether mutations in this site impact regulatory interactions in the -142 kb assay. Figure 4A and B show the control *G-nhomie-L* and the corresponding Su(Hw) binding site mutant transgene, *G-nhomieΔSH-L*. As evident from a comparison of wild-type and *Su(Hw)* mutant embryos, expression of the *LacZ* reporter both at the blastoderm stage and later in development is largely lost when the *nhomie* Su(Hw) binding site is mutated. Quantification of *LacZ* and *GFP* confirmed the staining results (Figure 4C).

**Figure 4.**
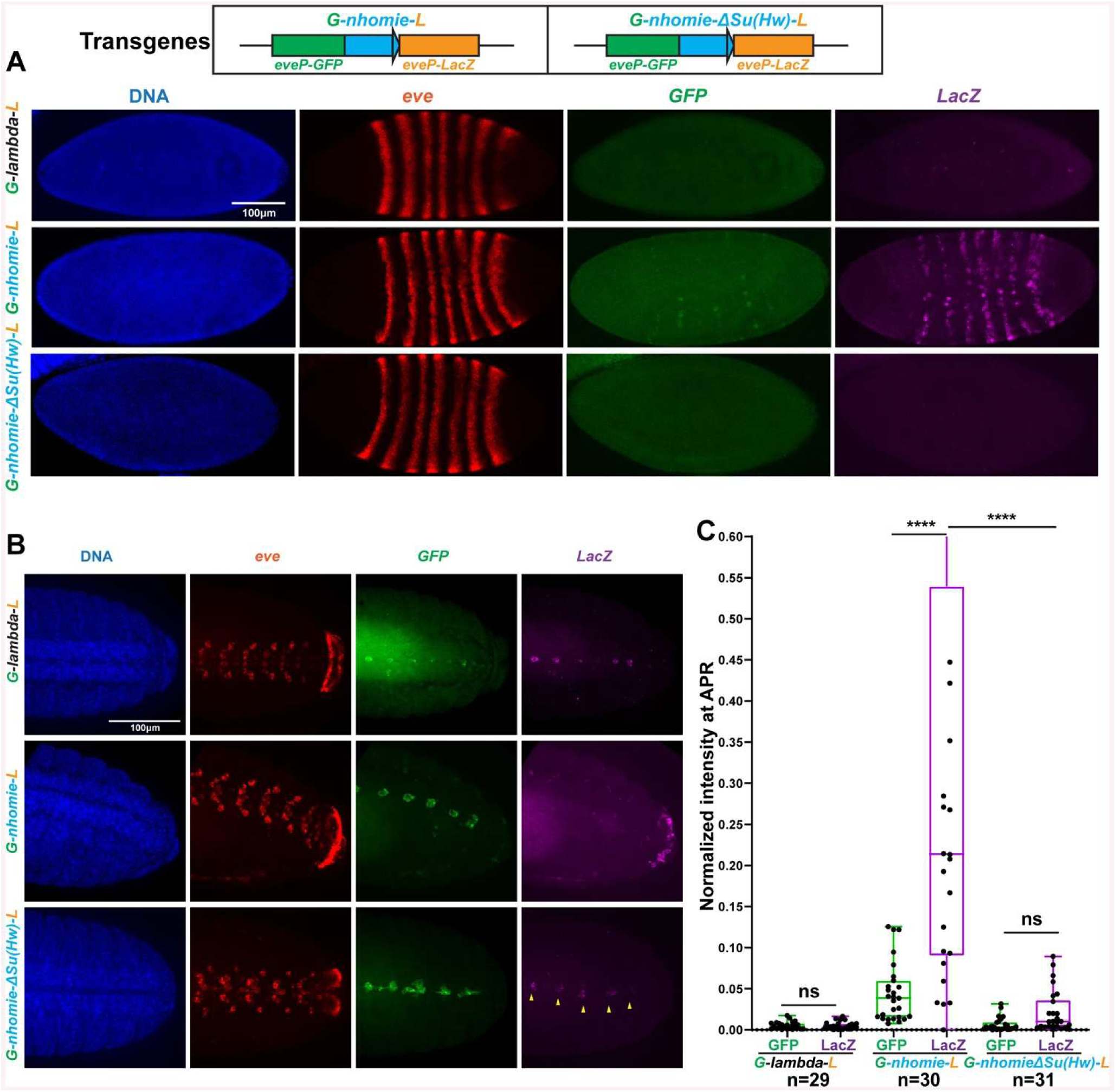
Mutation of *nhomie* Su(Hw) binding site disrupts activation of transgene reporter expression by the *eve* enhancers. A) The panels in each row show DAPI staining, *eve* mRNA, *GFP* mRNA, and *LacZ* mRNA for *G-lambda-L*, *G-nhomie-L*, and *G-nhomieΔSu(Hw)-L* in blastoderm stage embryos. B) The panels in each row show DAPI staining, *eve* mRNA, *GFP* mRNA, and *LacZ* mRNA for *G-lambda-L*, *G-nhomie-L*, and *G-nhomieΔsu(Hw)-L* in stage13-14 embryos. C) Normalized maximum intensity projections of *GFP* and *LacZ* mRNA staining in the APR of stage13-14 embryos for each transgene.

As the digoxigenin *in situ* hybridization procedure can be used to amplify weaker *LacZ* signals, we also used it to assay the effects of mutating the *nhomie* Su(Hw) binding sites, and this is shown in Figure 5A and B. While little if any *LacZ* expression was detected using HCR-FISH, *LacZ* transcripts driven by the *eve* enhancers can be detected in blastoderm and early gastrula *G-nhomieΔSH-L* embryos using digoxigenin *in situ*. The number of *LacZ*-positive cells is, however, substantially reduced compared to WT *nhomie*. Interestingly, in older germband extended and germband retracted embryos, *eve* enhancer-dependent expression in the APR and the CNS is largely absent. The loss of the Su(Hw) binding site also impacts the blocking activity of *nhomie*, since *LacZ* expression driven by the *hebe* enhancers in midline cell clusters is now observed (Figure 5B, stage 13, “13” and “13v”).

**Figure 5.**
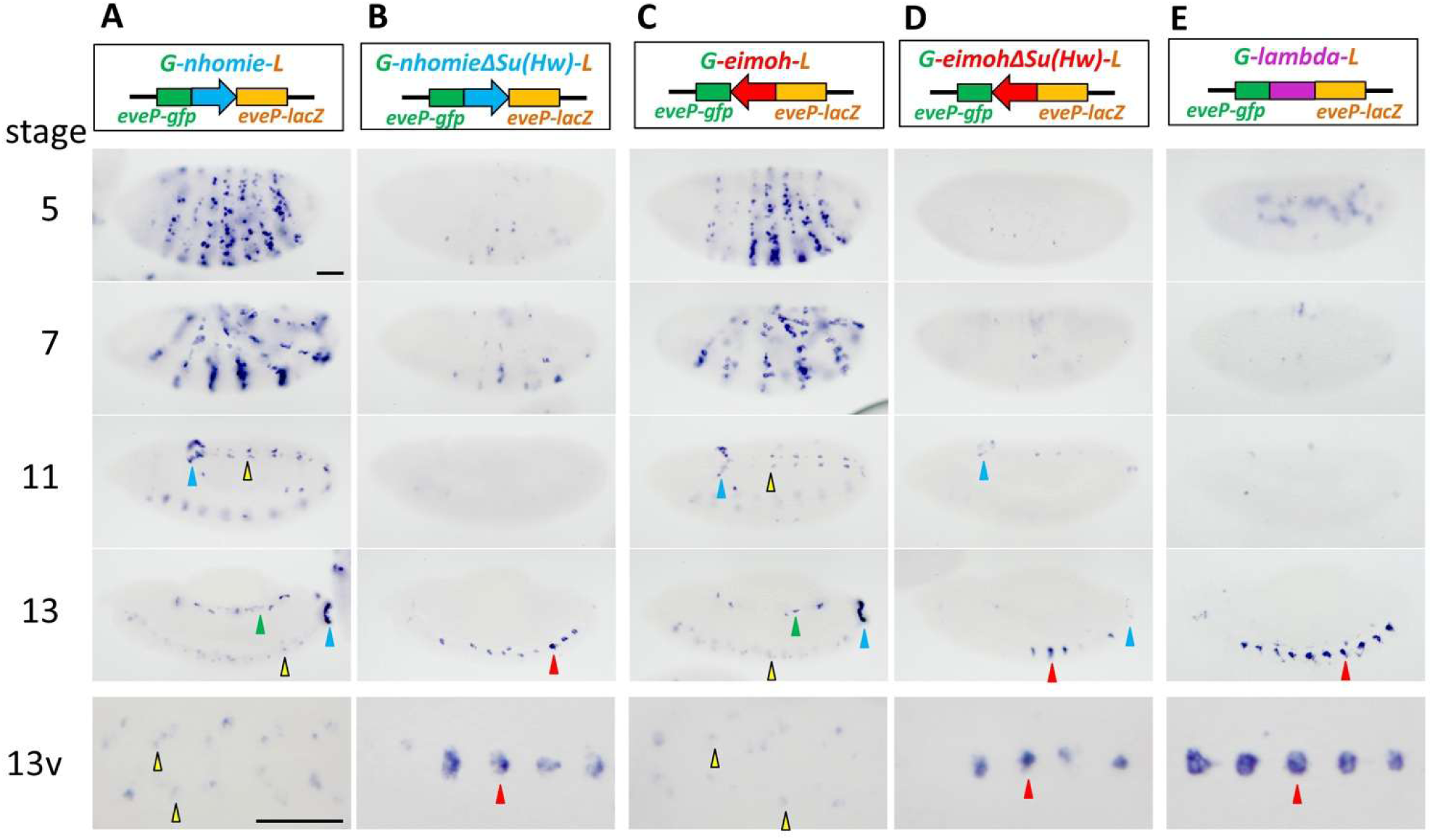
Digoxigenin *in situ* hybridization in wild type and the Su(Hw) binding site mutants in *nhomie* and *homie*. Expression of the *LacZ* reporter by A) *G-nhomie-L*, B) *G-nhomieΔSu(Hw)-L*, C) *G-eimoh-L*, D) *G-eimohΔSu(Hw)-L*, and E) *G-lambda-L* transgene inserts are shown. The transgene constructs are illustrated on the top. Embryonic stages 5, 7, 11, 13 are shown. Ventral views of stage 13 (13v) are shown in the bottom row. Arrowheads indicate the following: blue: APR, yellow: *eve*-expressing neuronal cells, green: *eve*-expressing mesodermal cells, and red: *hebe*-expressing midline cell clusters. Scale bar: 50μm.

### The Su(Hw) site in homieCDEF is required for long-distance regulatory interactions

We also reexamined the effects of Su(Hw) binding site mutation in *homie* on long-distance regulatory interactions. Instead of the 271 bp DEF *homie* element used in our previous study (Fujioka et al., 2025), we used a larger 367 bp *homie* element, CDEF. Like *nhomie*, the relative orientation of the *homie* in the transgene determines which of the two reporters is preferentially activated by the *eve* enhancers. For these experiments we oriented the wild-type *homie* and the Su(Hw) mutant *homieΔSH* in the transgene so that *LacZ* was located “upstream” of the transgene boundary (Figure 5C and D, and Figure 6; *G-eimoh-L* and *G-eimoh-ΔSu(Hw)-L*). In this configuration, the *LacZ* reporter is activated by the *eve* enhancers when the transgene *homie* pairs with the *eve* boundaries. The transgenes were then inserted into the -142 kb attP site so that the *LacZ* reporter is located on the *eve* side of the transgene *homie*.

**Figure 6.**
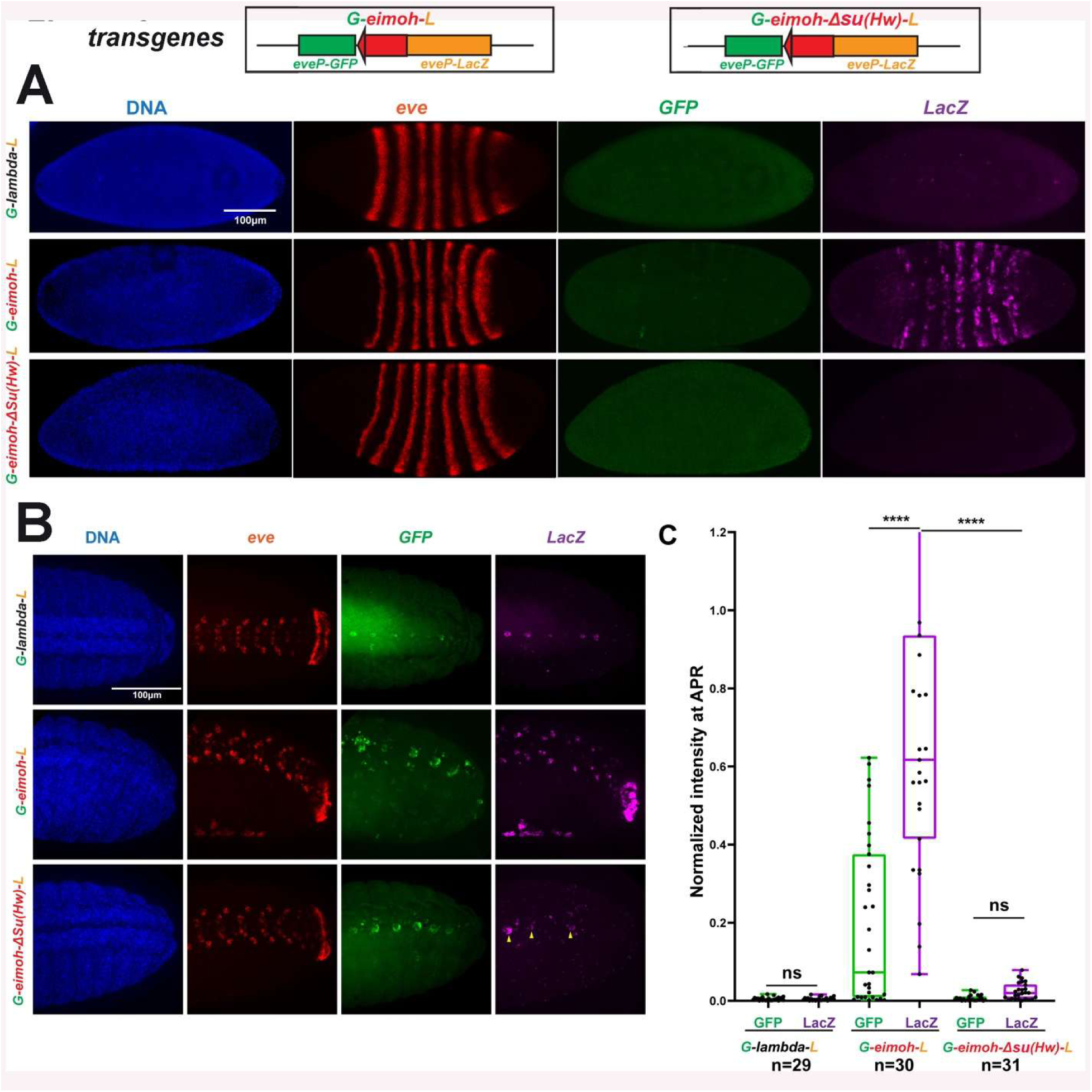
Mutation of the *homie* Su(Hw) binding site disrupts activation of transgene reporter expression by *eve* enhancers. **A)** The panels in each row show DAPI staining, *eve* mRNA, *GFP* mRNA, and *LacZ* mRNA for *G-lambda-L*, *G-eimoh-L*, and *G-eimohΔSu(Hw)-L* in blastoderm stage embryos. B) The panels in each row show DAPI staining, *eve* mRNA, *GFP* mRNA, and *LacZ* mRNA for *G-lambda-L*, *G-eimoh-L*, and *G-eimohΔSu(Hw)-L* in stage 13-14 embryos. C) Normalized maximum intensity projections of *GFP* and *LacZ* mRNA staining in the APR of stage 13-14 embryos for each transgene.

Figure 6 shows *G-eimohΔSH-L* HRC-FISH *in situs*, while Figure 5C and D shows digoxigenin *in situs*. Little or no *eve*-enhancer-dependent *LacZ* expression can be detected in *G-eimohΔSH-L* in stage 5 (or 7) embryos using either *in situ* hybridization procedure. Thus, the loss of the Su(Hw) binding site appears to have even greater impact on *homie-*dependent regulatory interactions at this stage in development than observed for *nhomie* (compare Figure 5B and D). On the other hand, in stage 11 and 13 embryos in *G-eimohΔSu(Hw)-L*, weak *LacZ* expression can be detected in the APR, but not in *G-nhomieΔSu(Hw)-L.* Quantification of *LacZ* and *GFP* expression in the APR confirmed the staining results (Figure 6C). For both cases, several midline cell clusters in the CNS were observed, indicating that in addition to being largely unable to mediate long-distance regulatory interactions, the Δ*SH* mutant is unable to fully block the *hebe* enhancers from activating *LacZ* expression (Figure 4B, 5B, 5D and 6B).

### Mutation of Su(Hw) binding sites in nhomie and homieCDEF disrupts long-distance physical interactions

To better understand how the loss of the Su(Hw) binding sites in *nhomie* and *homie* impacts the ability of these TAD boundaries to mediate long-distance regulatory interactions, we used MicroC to probe the physical interactions between the transgenes and the *eve* TAD. The MicroC contact maps for *G-nhomie-L* and *G-nhomieΔSH-L* are shown in Figure 7A and B, while Figure 7C and D show “virtual 4C” viewpoints from the transgene *LacZ* and *GFP* reporters (similar to the analysis in Figure 3). As expected from the substantial reduction in reporter activity, contacts between the *LacZ* reporter in *G-nhomieΔSH-L* and sequences in the *eve* TAD are diminished. This change can be seen in the blow-ups in panels 7A and B and in the *LacZ* and *GFP* viewpoints in panels C and D. Quantitation indicates that there is a nearly 10-fold reduction in contact frequency (Figure 7E). Interestingly, however, the pattern of physical interactions between *G-nhomieΔSH-L* and the surrounding TADs, with *eve* and the TADs surrounding *eve* appear to be quite similar to that of the wild-type *G-nhomie-L* insert (Figure 7-figure supplemental 1). This would suggest that the transgene boundary still pairs in the same orientation-dependent manner with the endogenous *eve* locus, just less frequently or less stably than it does when the Su(Hw) site is intact.

**Figure 7.**
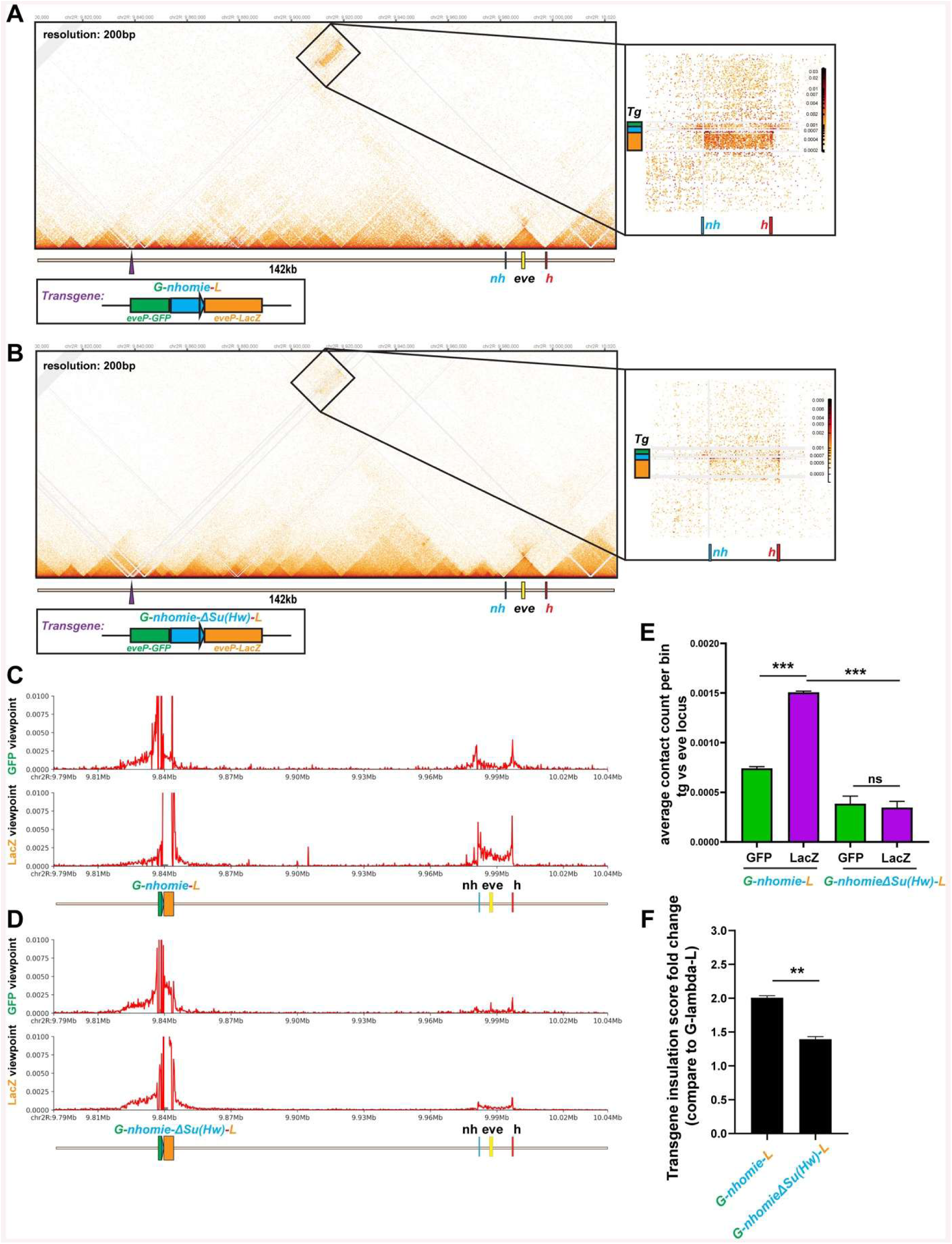
Mutation of the *nhomie* Su(Hw) binding site disrupts physical interactions between the transgene and sequences in the *eve* TAD. The MicroC contact profile and blow up for A) *G-nhomie-L* and B) *G-nhomieΔSu(Hw)-L* transgenes. Viewpoints from the *GFP* and *LacZ* reporters in C) *G-nhomie-L* and D) *G-nhomieΔSu(Hw)-L* transgenes. E) Average contact count per bin for the *GFP* and *LacZ* reporters, as indicated for the two transgene inserts *G-nhomie-L* and *G-nhomieΔSu(Hw)-L*. F) Insulation score for the transgene boundaries *nhomie* and *nhomieΔSu(Hw)*.

The MicroC contact profile for the *G-eimoh-L* and *G-eimohΔSH-L* inserts are shown in Figure 8A and B. As was the case for the *G-nhomieΔSH-L* insertion, there is a substantial reduction in the physical interactions between the *G-eimohΔSH-L* and sequences in the *eve* TAD. This can be seen by comparing the *LacZ* and *GFP* reporter “virtual 4C” viewpoints for *G-eimoh-L* in Figure 8C with those for *G-eimohΔSH-L* in Figure 8D. Even though the frequency of physical interactions is greatly reduced (Figure 8E), the residual pattern of interactions between *G-eimohΔSH-L* and the surrounding TADs with *eve* and the TADs surrounding *eve* still resemble that seen for *G-eimoh-L*. Thus, like *nhomieΔSH*, the *eimohΔSH* retains some orientation-dependent pairing activity.

**Figure 8.**
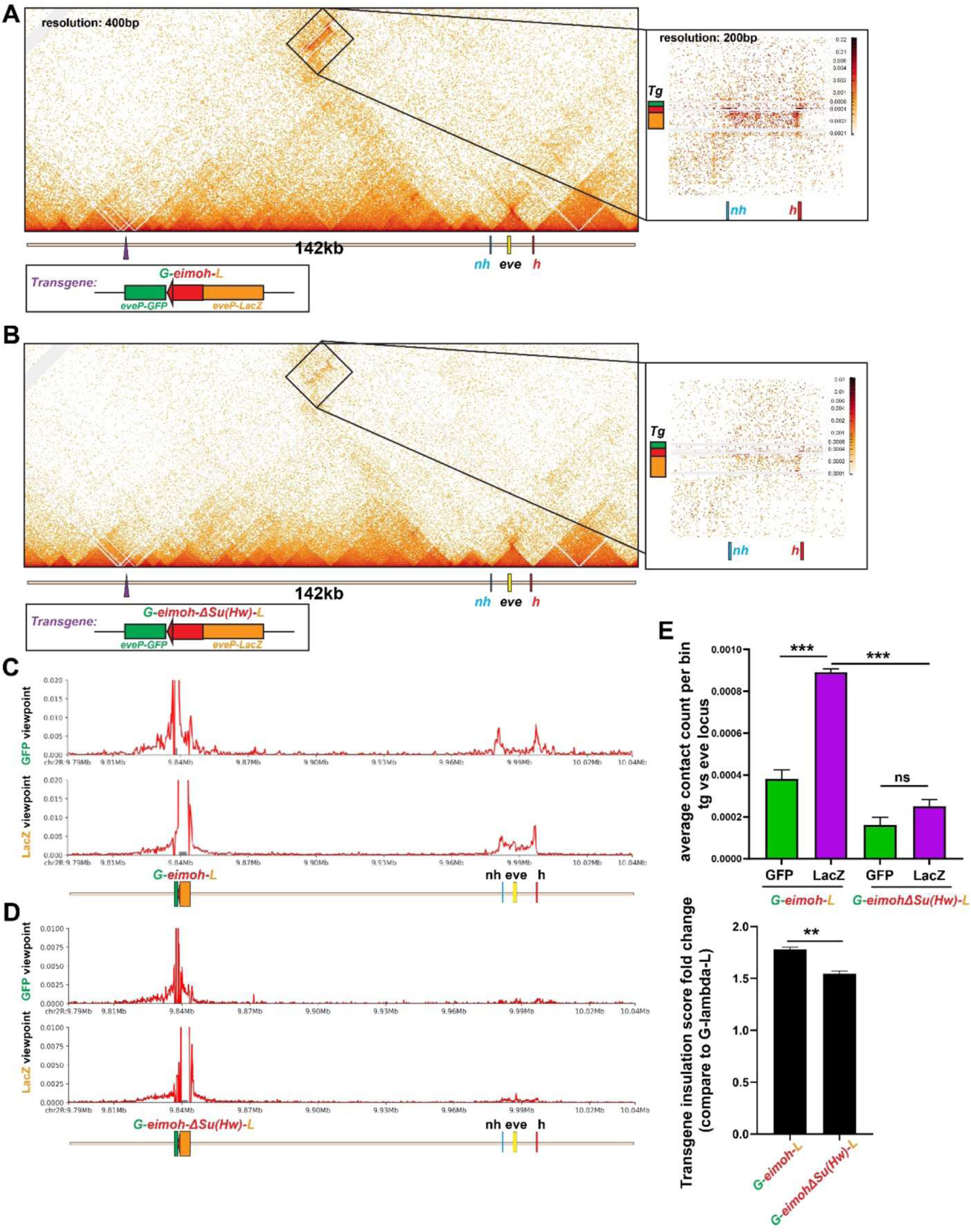
Mutation of the *homie* Su(Hw) binding site disrupts physical interactions between the transgene and sequences in the *eve* TAD. MicroC contact profile and blow up for A) *G-eimoh-L* and B) *G-eimohΔSu(Hw)-L* transgenes. Viewpoints from the *GFP* and *LacZ* reporters in C) *G-eimoh-L* and D) *G-eimohΔSu(Hw)-L* transgenes. E) Average contact count per bin for the *GFP* and *LacZ* reporters, as indicated for the two transgene inserts *G-eimoh-L* and *G-eimohΔSu(Hw)-L*. F) Insulation score for the transgene boundaries *eimoh* and *eimohΔSu(Hw)*.

Since mutating the Su(Hw) binding sites in both *nhomie* and *homie* results in the activation of the *LacZ* reporter by the *hebe* enhancers located just beyond the *GFP* reporter, we calculated the “insulation” score. We used Fan-C and a window of 4 kb to analyze cross-TAD contacts for *lambda* DNA, the wild-type and Su(Hw) mutant *nhomie* and *homie* boundaries (see Methods). For this purpose, we measured the contact frequency between sequences immediately upstream and downstream of the transgene inserts. Figure 7F shows the insulation score for wild -type and Su(Hw)-mutant *nhomie* relative to that of *lambda*, while Figure 8F shows the insulation score for wild-type and Su(Hw)-mutant *homie* relative to that of *lambda*. In both cases, there is a significant drop due to the Su(Hw) binding site mutation.

### Long-distance physical interactions between the transgene boundary and the endogenous nhomie and homie

The regulatory interactions between the reporters in the transgene insert and the *eve* enhancers depend upon the physical pairing of the transgene boundary with the endogenous *nhomie* and *homie*. In previous experiments (Bing et al., 2024), we found that when the viewpoint is centered on *homie* in the transgene rather than one of the reporters, the interactions with *eve* peak at the *nhomie* and *homie* boundaries (Figure 9C). This is also true for transgenes containing the *nhomie* boundary. Figure 9A shows that the interactions between *nhomie* in the *G-nhomie-L* transgene and *eve* are centered on the *nhomie* and *homie* boundaries. These findings support the conclusion that pairing of the transgene boundary with the two *eve* boundaries is directly responsible for physically linking the transgene to the *eve* TAD.

**Figure 9.**
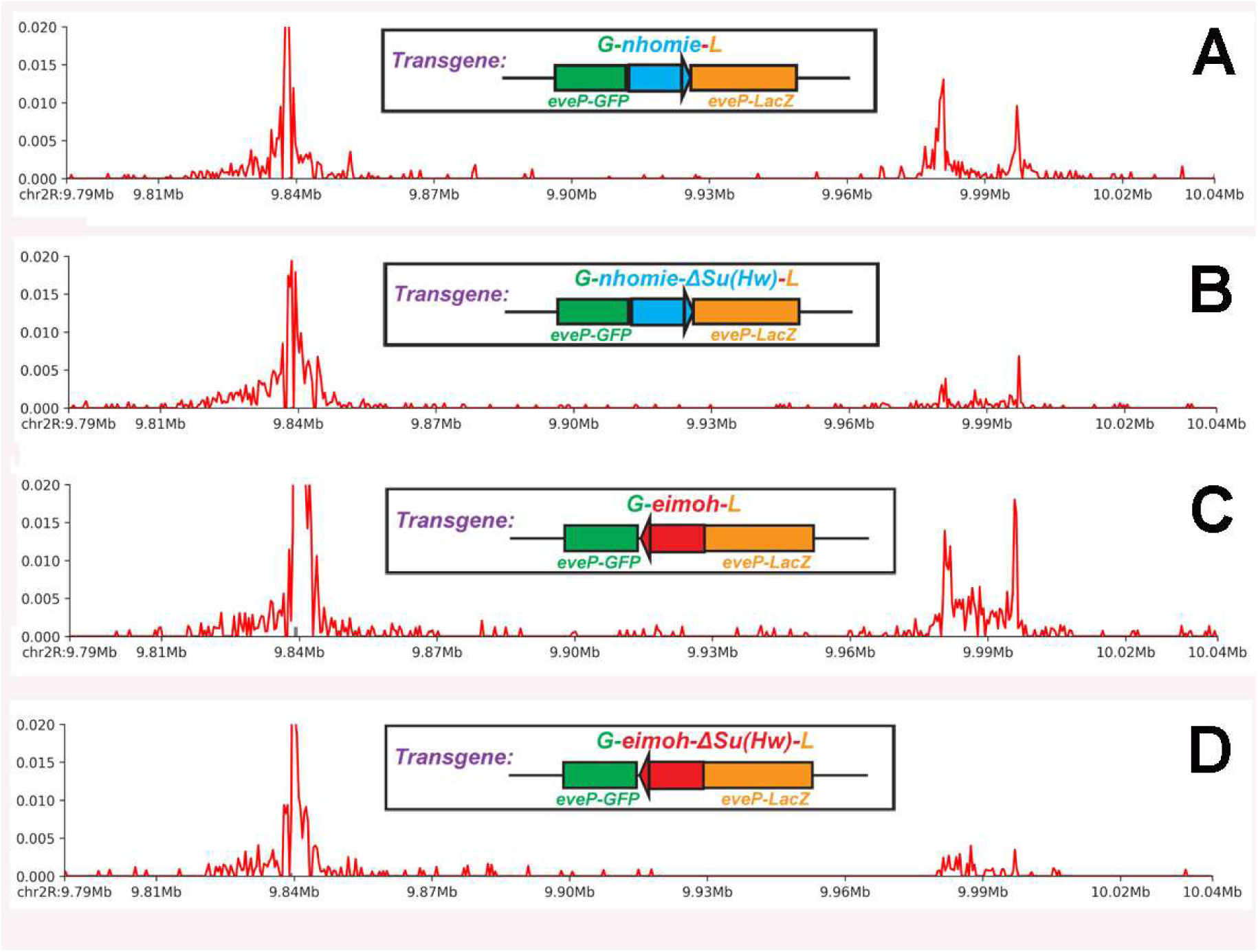
Viewpoints from the transgene boundaries. A) *nhomie*, B) *nhomieΔSu(Hw)*, C) *eimoh*, and D) *eimohΔSu(Hw)*. Note the reduction in contacts between the Su(Hw) site mutant boundaries in the transgene and both *nhomie* and *homie* (at the ends of the *eve* TAD).

As would be predicted by the boundary pairing model, mutations in the Su(Hw) binding site result in a disruption of the physical interactions between the transgene boundary and *nhomie* and *homie* in the *eve* locus. For the Su(Hw) mutation in *G-nhomieΔSH-L*, the physical interactions between the transgene boundary and the two *eve* boundaries are substantially reduced (Figure 9B). Moreover, consistent with the idea that the shared Su(Hw) binding sites are important for *nhomie* self-pairing and for heterologous pairing with *homie*, the prominent peaks for *nhomie*-transgene with both *nhomie* and *homie* are substantially reduced in the *nhomieΔSH* viewpoint (Figure 9B). The physical interactions are not completely lost, as there are still small peaks that map to both *nhomie* and *homie*.

The physical contacts between the transgene *homieΔSH* and the two *eve* boundaries are also compromised by the Su(Hw) binding site mutation (Figure 9C and D). Moreover, as was the case for *nhomie*, *homie* self-pairing (*homie*-transgene with *homie*) and heterologous pairing (*homie-*transgene with *nhomie*) largely depend upon the shared Su(Hw) binding sites.

### *homie* and *nhomie* Su(Hw) mutant boundaries retains] pairing activity in transvection assays

While the Su(Hw) mutations disrupt long-distance interactions between the transgene boundary and the boundaries in the *eve* TAD, weak physical interactions can still be detected in viewpoints from the reporters and the transgene boundary. This would suggest that factors other than Su(Hw) must be involved in pairing interactions, and that they might be sufficient on their own to mediate physical interactions in less demanding assays.

To test this possibility, we used a transvection assay in which a *LacZ* reporter transgene interacts with a transgene containing enhancers, each inserted into an *attP* site distant from *eve*, where long-range interactions with the endogenous *eve* enhancers are not observed (Fujioka et al., 2016). As pairing interactions in *trans* in this *attP* environment are relatively weak, the reporter is only weakly activated by the *eve* enhancers when the *LacZ* reporter carries *lambda* DNA and the enhancer transgene carries the *homieCDEF* that was used in -142kb assay (Figure 10A). In contrast, when both transgenes have the *homieCDEF* inserted in the *same* orientation relative to the reporter and the enhancers, the *LacZ* reporter is activated strongly by the *eve* mesodermal and APR enhancers (Figure 10B). *Trans-*activation is also observed when the 367 bp *homie* fragment CDEF is paired with the smaller 271 bp *homie* fragment DEF (Figure 10C). In contrast to the -142 kb assay, the regulatory interactions in the transvection assay are less sensitive to mutations in the Su(Hw) binding site. While mutation of the Su(Hw) site in DEF (DΔSuEF, located in the reporter transgene) weakens regulatory interactions, the *eve* enhancers are still able to active *LacZ* expression more strongly than in the *lambda* DNA control (Figure 10D relative to 10A).

**Figure 10.**
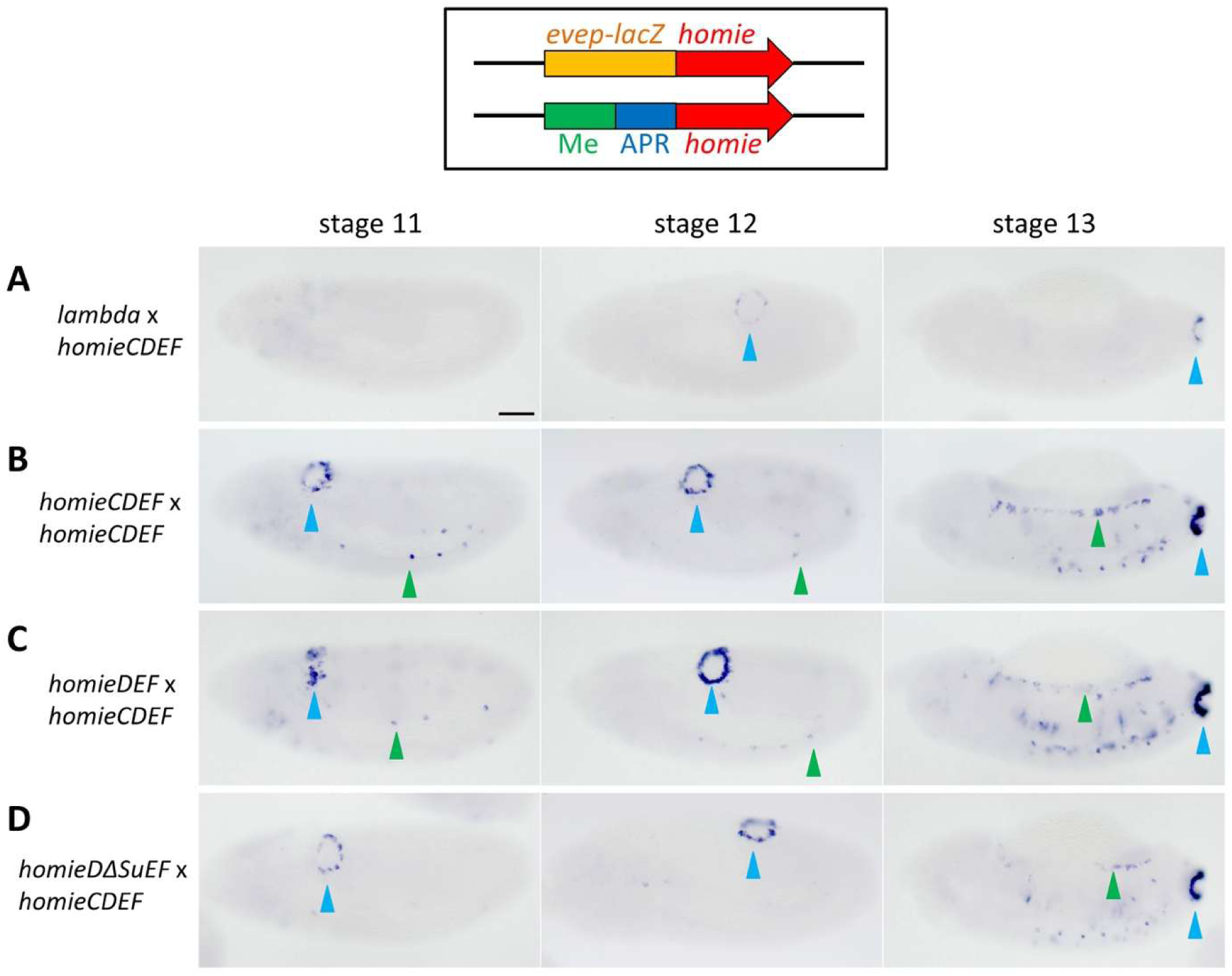
*homie*-dependent transvection is weakened by loss of the Su(Hw) binding site. Digoxigenin *in situ* hybridization showing *LacZ* expression in the transvection assay. The two transgene constructs used to assay transvection are illustrated at the top. The different transgene constructs used for each cross are indicated on the left as: reporter x enhancer. The negative control has a 500 bp *lambda* DNA fragment. The 367 bp *homie* (*homieCDEF*), 271 bp *homie* (*homieDEF*), and the Su(Hw) site-mutated *homieDEF* (*DΔSuEF*) were tested. Stages 11, 12, and 13 are shown. Arrowheads indicate the following: blue: APR and green: mesoderm. Scale bar: 50μm.

We performed a similar set of transvection experiments with *nhomie*. However, instead of using the full length 600 bp *nhomie* fragment we used a truncated 200 bp fragment. Importantly, the long-distance pairing activity of the truncated *nhomie* fragment is substantially reduced compared to the 600 bp fragment (Park, 2026). Despite its reduced pairing activity in the long-distance assay, it still functions in the transvection assay (Figure 10 figure supplemental 1). Moreover, mutations in the Su(Hw) site in the 200 bp fragment weakened but do not eliminate stimulation by the *eve* enhancers.

### Shared Su(Hw) binding sites promote pairing interactions in the transvection assay

In previous studies (Fujioka et al. 2025), we found that the *gypsy* insulator is unable to mediate regulatory interactions between transgenes inserted at -142 kb and the *eve* enhancers (Figure 11**-**figure supplemental 1B). One interpretation of this finding is that the Su(Hw) proteins associated with *homie* or *nhomie* are not in themselves sufficient for physical interactions with the *gypsy* insulator. An alternative possibility is that while the shared Su(Hw) proteins in *nhomie/homie* can engage in physical interactions with the *gypsy* insulator these interactions are not strong enough to mediate the sustain physical contacts needed to activate reporter transcription over a distance spanning 142 kb and multiple intervening TADs.

**Figure 11.**
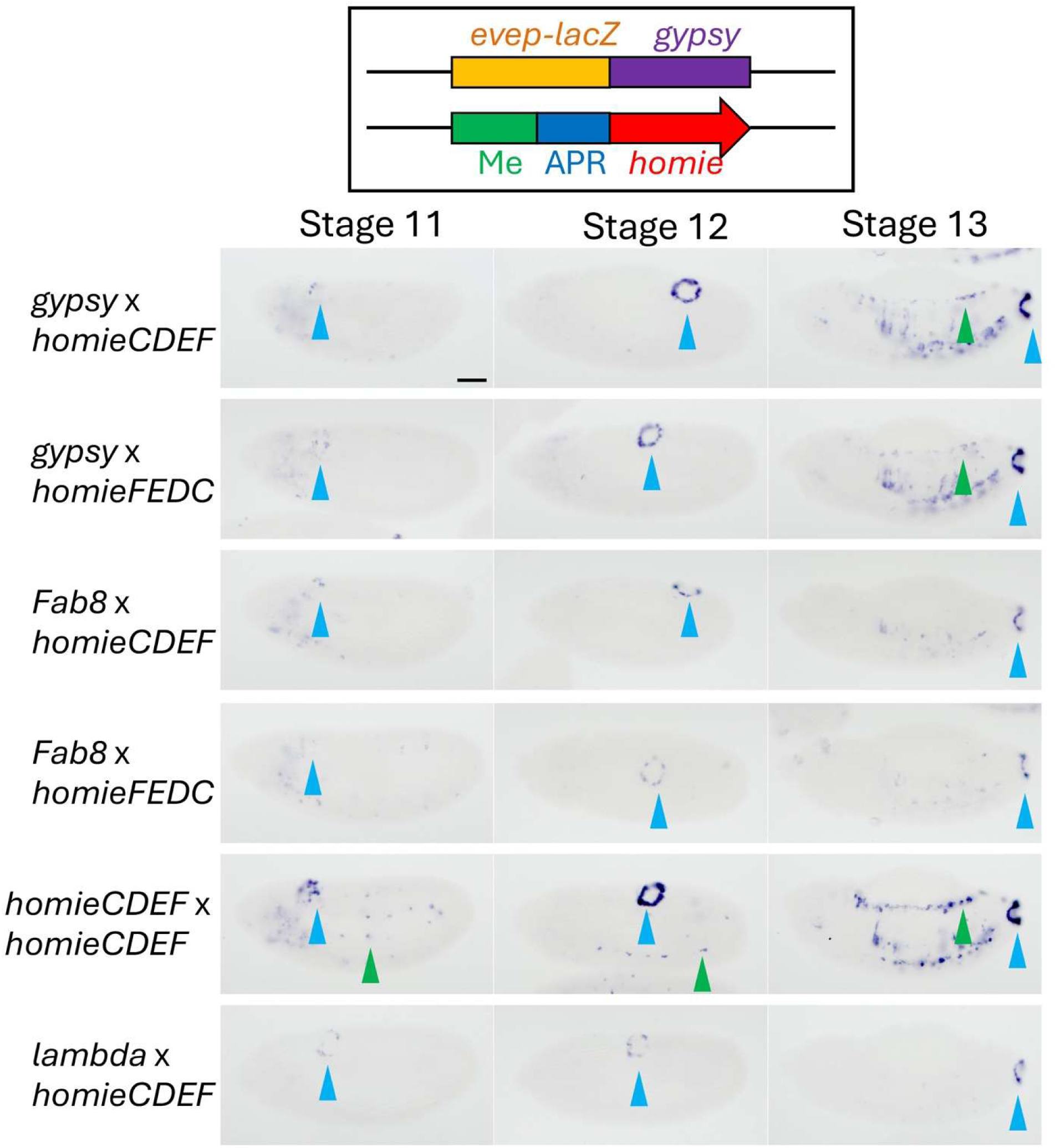
The su(Hw) *gypsy* insulator supports transvection with *homie*. Digoxigenin *in situ* hybridization showing *LacZ* expression in the transvection assay. The two transgene constructs for each experiment are shown on the top. The transgene combinations used in each case are indicated as reporter x enhancer. The negative control has a 500 bp *lambda* DNA fragment. The 349 bp *gypsy* fragment, the 367 bp *homie* (*homieCDEF* and *homieFEDC*) fragments and the 329 bp *Fab-8* fragment (Kyrchanova et al., 2016) were tested. Note that the orientation of *homie* in *homieFEDC* is inverted in the transgene to test whether *gypsy* interaction with *homie* is orientation-dependent. Stages 11, 12, and 13 are shown. Arrowheads indicate the following: blue: APR and green: mesoderm. Scale bar: 50μm.

To distinguish between these possibilities, we used the transvection assay to test whether the single Su(Hw) binding site in *homie* is sufficient to mediate regulatory interactions with the *gypsy* insulator. Figure 11 shows that the interactions between *homieCDEF* and the *gypsy* insulator in this transvection assay are not altogether different from when *homie* is paired with itself. Moreover, unlike *homieCDEF:homieCDEF* pairing, *gypsy:homieCDEF* pairing is orientation independent: the *eve* enhancers can activate *LacZ* expression independent of the relative orientation of the *gypsy* and *homieCDEF* elements at a level only slightly lower than that observed for *homieCDEF:homieCDEF* pairing interactions. Although we did not test orientation dependence for *gypsy:nhomie* pairing, the *gypsy*-containing reporter is activated by the *eve* enhancers when the enhancer transgene has *nhomie* (Figure 11-figure supplemental 1A).

These results show that the two *eve* boundaries can productively partner with the *gypsy* insulator in a less demanding assay. This is not unexpected as fly boundaries are known to be somewhat promiscuous in their pairing interactions (c,f, Kychanova et al., 2011; Gohl et al., 2011; Li et al., 2018). In this case, the likely reason would be the shared Su(Hw) binding sites. Consistent with this idea, we found that the dCTCF dependent *Fab-8* boundary (Moon et al., 2005; Kyrchanova et al., 2016) does not partner with *homieCDEF* in the transvection assay (Figure 11).

## Discussion

TAD formation in the boundary:boundary pairing model depends upon physical interactions between proteins that are associated with interacting boundary elements. In one mechanism, boundaries that share binding sites for the same DNA binding protein can be linked together if that protein can assemble into homodimers and/or homomultimers. This mechanism is likely used by polydactyl zinc finger proteins like CTCF, Zw5, Zipic, and Pita, by the BEN domain protein Insensitive, and DNA binding proteins like GAF, Mod(mdg4), and CP190 that have BTB domains (Fedolova et al., 2017; Bonchuck et al., 2021). Unlike these chromosomal architectural proteins, Su(Hw) is not known to form homomeric complexes; however, previous studies have suggested that its chromosome architectural functions depend upon shared binding sites. The first evidence for this possibility came from studies by Sigrest and Pirrotta (1997), who showed that transgenes carrying the Su(Hw)-dependent *gypsy* insulator and a Polycomb response element from the Bithorax complex could mediate long-distance Polycomb-dependent silencing of a *white* reporter. Subsequent work by Cai and Shen (2001) and Muravyova et al. (2001) presented additional genetic evidence for the pairing of *gyspy* insulators, and this was extended by the boundary bypass experiments of Kyrchanova et al. (2008a). In these experiments, Kyrchanova et al. showed that bypass depended on shared binding sites for chromosomal architectural proteins. When multimerized binding sites for Su(Hw), CTCF, or Zw5 were paired with themselves, bypass was observed; however, when multimerized Su(Hw) binding sites were paired with multimerized CTCF or Zw5 binding sites, there was no bypass. The Kyrchanova et al. paper also showed that self-pairing is a common property of endogenous boundaries in flies.

While the genetic experiments showing that fly boundaries utilize shared binding sites in their pairing interactions are compelling, a direct demonstration of the physical interactions that are mediated by these shared sites is lacking. In fact, there is limited evidence linking binding sites for fly chromosomal architectural proteins with the physical interactions involved in self or heterologous boundary:boundary pairing. To address this question, we have used the *eve* TAD boundaries, *nhomie* and *homie*, to test the role of shared binding sites. Genetic studies have shown that, like other fly boundaries, *nhomie* and *homie* pair with themselves head-to-head. They also pair with each other, in this case in a head-to-tail configuration. This pairing interaction generates a TAD with a stem-loop topology, which has a distinctive contact profile in MicroC experiments (Bing et al. 2024; Ke et al. 2024). When *nhomie* or *homie* is included in a dual reporter transgene inserted in an *attP* site 142 kb from *eve*, the *eve* enhancers drive reporter expression. In previous studies on *homie*, we showed that these regulatory interactions accompany the physical pairing of the transgene boundary with both of the endogenous *eve* boundaries (Bing et al., 2024). As reported here, this is also true for *nhomie* (Figs. 2 and 3). This can be seen when the transgene *nhomie* is used as a viewpoint: it contacts both *nhomie* and *homie* in the *eve* locus. Also like *homie*, the relative orientation of *nhomie* in the transgene determines which of the two reporters is activated. Moreover, the pattern of activation is a reflection of the physical contacts that are visualized with MicroC between the two reporters and sequences in the *eve* TAD.

### Mutations in the shared homie and nhomie Su(Hw) binding sites disrupt self and heterologous pairing interactions

As would be predicted if shared sites for DNA binding proteins are important for boundary function, mutations in the *homie* and *nhomie* Su(Hw) sites largely abrogate long-distance regulation by the *eve* enhancers (Figure 4, 5, and 6). Although the effects of the Su(Hw) mutation on transcriptional activation are in both cases substantial, we are able to detect regulatory interactions using the more sensitive digoxigenin *in situ* procedure (Figure 5). For *nhomieΔSH*, we detect residual *LacZ* expression in an *eve-*like stripe pattern at the blastoderm and early gastrulate stages, while there is little if any expression evident in stage 11 and 13 embryos, when *eve* is expressed in the APR, CNS, and mesoderm. In contrast to *nhomieΔSH*, only very few *LacZ*-positive cells are visible in blastoderm and early gastrula embryos in *eimohΔSH*. Instead, we detect weak *LacZ* expression in stage 11 and 13 embryos in the APR and CNS. It is not clear why enhancer-driven *LacZ* expression for *nhomieΔSH* seems to be restricted to blastoderm and early gastrulation embryos, while for *eimohΔSH*, enhancer-dependent expression is stronger in stage 11 and 13. One possibility is that residual self-pairing of the Su(Hw) mutant boundary is stronger than pairing with the heterologous boundary. This is suggested by the proximity of the relevant enhancers to the *eve* boundary whose transgene partner carries the Su(Hw) mutation. The 7-stripe enhancer that drives *eve* expression during early gastrulation is located next to the *nhomie* boundary, and this is the time when *nhomieΔSH-*dependent *LacZ* expression is highest. Likewise, the APR and mesodermal enhancers are located between *eve* and *homie*, and thus would be closer to the *homie* boundary. Another possibility is that *nhomie* and *homie* have different stage/tissue-specific boundary factors: the *nhomie* stage specific factors would be present in early embryos, while the *homie* stage/tissue specific factors would be present later in embryogenesis.

### The homie Su(Hw) binding site mutant retains some boundary function

Although the long-distance regulatory and physical interactions of *GeimohΔSHL* with the *eve* locus is significantly reduced, it is not completely disrupted. Consistent with the idea that the mutant retains some boundary activity, we can readily detect *LacZ* expression driven by *eve* enhancers when a minimal 271 bp *homie* element, *DΔSuEF*, carrying the Su(Hw) binding site mutation is combined with the standard 367 bp (CDEF) *homie* boundary in our transvection assay (Figure 10). The likely reason is that this is a less demanding assay, as boundaries to either side of the reporter and enhancer transgenes in the chromosome are expected to be mediating homologous pairing (via head-to-head self-pairing interactions), and this would help align boundaries in the transgene and stabilize their physical interactions. By contrast, the -142 kb assay boundary pairing takes place across multiple TADs, and, as is evident from the novel contacts that are induced, requires a substantial distortion in the native organization of the chromatin fiber in between the transgene and the *eve* TAD, as well as to either side. This would likely tend to disfavor physical interactions between the distant boundary elements.

### Shared binding sites and TAD formation

Taken together our findings support the idea that one of the mechanisms orchestrating the physical pairing of TAD boundaries in *cis* is the presence of shared binding sites for the same chromosomal architectural proteins. Consistent with this idea, recent computational analysis of the TAD boundary lexicon in *Drosophila* identified 575 TAD boundaries in which Su(Hw) is predicted to play a key role in TAD formation (Wang et al., 2026). Of these, 138 correspond to adjacent boundaries that like *nhomie* and *homie* share Su(Hw) binding sites and define the endpoints of a TAD. On the other hand, most of the predicted Su(Hw) dependent TAD boundaries are solo—namely, neither of its neighbors has a Su(Hw) site that is predicted to be functionally important. If these predictions are correct, the Su(Hw) protein in the solo TAD boundary would have to partner with some other protein(s) that is bound to one (or both) of the neighboring boundaries. Clearly it will be important to identify factors that can engage in heterologous interactions of this type.

### Specificity versus promiscuity in pairing interactions

Although there are multiple TAD boundaries between the -142 kb attP site and *eve*, none have Su(Hw) binding sites, and there is no indication of pairing interactions between these boundaries and either *homie* or *nhomie* in the MicroC contact profiles and the viewpoints from the transgene towards *eve* (Figure 2, 3, 7, 8, and 9). ChIP experiments indicate that there are two Su(Hw) peaks to the left of the -142 kb attP site that could potentially interact with the transgene boundaries or the endogenous *eve* boundaries (Figure 7-figure supplemental 2). However, we did not detect any pairing interactions in the MicroC contact profiles. This would suggest that the presence of a set of shared binding sites in two TAD boundaries in the neighborhood is not in itself sufficient for cross-TAD interactions. Consistent with this idea, we found that a 451 bp *gypsy* transposon Su(Hw) element was unable to mediate regulatory interactions between a transgene reporter inserted at -142 kb and the *eve* enhancers (Fujioka et al. 2025: see Figure 11-figure supplemental 1B). On the other hand, shared Su(Hw) sites would appear to be sufficient for pairing interactions in less demanding assays, as the *gypsy* insulator is able to support transvection when paired with *homie* (Figure 11) or *nhomie* (Figure 11-figure supplemental 1A). In both cases, the level of reporter expression is much closer to that observed when *homie* or *nhomie* are paired with themselves than it is to that seen when one of the transgenes carries *lambda* DNA in place of a boundary. Thus, the observed levels of specificity versus promiscuity depend upon the design of the assays that are used to detect pairing interactions. Interestingly, even in the less demanding transvection assay some degree of specificity is retained as the dCTCF dependent *Fab-8* boundary is unable to partner with *homie*.

### Mechanisms underpinning TAD assembly

Two different mechanism have been proposed to explain how TADs are formed. One is the physical pairing of neighboring boundary elements in *cis* that has been described here, while the other is the cohesin loop extrusion/CTCF road block model (Dixon et al., 2012; Rao et al., 2017; Fudenberg et al., 2017; Davidson and Peters, 2021; Dekker and Mirny, 2024). The results reported here together with those presented in a previous publication on the pairing properties of the other *eve* boundary, *homie*, cannot be explained by the cohesin loop extrusion model; however, they are completely consistent with a mechanism for TAD formation that depends on the physical pairing of compatible boundary elements.

In order to activate reporters inserted in the -142 kb attP site the *eve* enhancers need to be brought into close proximity. In the loop extrusion model, the cohesin complex would have to break through multiple intervening TAD boundaries and come to a halt when it encounters the transgene *nhomie* or *homie* boundary. This would generate a stem-loop in which cohesin sits atop the *eve homie* boundary and the transgene boundary and holds them together. In this configuration the reporter on the *eve* side of the transgene boundary will be in the same stem-loop as the *eve* enhancers and it will be activated by the *eve* enhancers. At the same time, the transgene boundary will block the *eve* enhancers from activating the reporter on the opposite, upstream side of the transgene boundary. If the *homie* and *nhomie* boundaries function as cohesin roadblocks, independent of their relative orientation with respecting to the extruding cohesin complex, the reporter on the *eve* side of the transgene boundary will always be activated, while the reporter on the opposite side of the transgene boundary will not.

In this paper, we tested the four possible *nhomie* transgene configurations: a) *nhomie* “pointing towards” *LacZ*; b) *nhomie* “pointing towards” *GFP*; c) *LacZ* on the *eve* side of the transgene boundary and d) *GFP* on the *eve* side of the transgene boundary. The results for two of the configurations—*LacZ* on the *eve* side of the transgene boundary *with nhomie* “pointing towards” *LacZ* (*G-nhomie-L*) and *GFP* on the *eve* side of the transgene boundary *with nhomie* “pointing towards” *GFP* (*L-nhomie-G*) are consistent with the predictions of the loop extrusion model. However, if one flips, for example, the orientation of the *G-nhomie-L* transgene in the attP insertion site so that the *GFP* reporter is on the *eve* side of the boundary (*L-eimohn-G*), the *GFP* reporter is not activated as would be predicted by the cohesin loop extrusion model. Instead, the *LacZ* reporter is activated by the *eve* enhancers. Since the orientation of the CTCF sites relative to the incoming cohesin complex is thought to determine whether the CTCF protein functions as a cohesin roadblock, it is possible that the inverted *nhomie* boundary is inactive as it failed to halt the extruding cohesin complex. However, this possibility is not correct as the inverted boundary blocks the *hebe* enhancers from turning on the *GFP* reporter. Essentially the same results are observed when the *nhomie* boundary in the *G-nhomie-L* insert is inverted to give *G-eimohn-L*. Even though the *LacZ* reporter is still on the *eve* side of the transgene boundary, it is not activated by the *eve* enhancers as would be predicted by the loop extrusion model. Instead, the *GFP* reporter is turned on by both the *eve* and *hebe* enhancers.

The shortcomings of the loop extrusion model are also clearly evident in the patterns of physical contacts seen in MicroC experiments. As predicted by the boundary pairing model, physical interactions between the two reporters and sequences in the *eve* TAD track completely with the relative reporter activity (Figs. 2 and 3 for *nhomie*; Figs. 8 and 9, see also Bing et al. 2024 for *homie*). Moreover, independent of the orientation of the boundary in the transgene, it physically interacts with both *nhomie* and *homie* in the *eve* TAD (see also Bing et al. 2024). The physical interactions generated when the *nhomie* boundary is reversed in *G-eimohn-LacZ* indicate that the ability to pair with boundaries in the *eve* TAD is independent of the orientation of the transgene boundary. This would rule out a model in which cohesin bypasses the “inappropriately” oriented *nhomie* or *homie* boundaries and stops at some imaginary TAD boundary beyond the *hebe* gene so that the reporter on the “wrong side” of the boundary is activated by the *eve* enhancers. In fact, there is no evidence of novel contacts between the boundaries in the *eve* TAD and boundaries beyond the transgene insert as would be expected if cohesin fail to stop when the transgene boundary is “inappropriately” oriented.

These are not the only shortcomings of the loop extrusion model. There is no mechanism in this model that could explain how TAD boundaries are able to promote *trans* regulatory interactions. However, we have shown here (Figure 10 and Figure 11-figure supplemental 1) and elsewhere (Fujioka et al., 2016) that *nhomie* and *homie* facilitate transvection either by pairing with themselves in *trans* or by pairing with each other in *trans*. For self-pairing, the *nhomie* or *homie* elements in the enhancer and reporter transgenes must be in the same orientation, while for heterologous pairing (*nhomie:homie*) they must be in the opposite orientation (Fujioka et al. 2016). Moreover, although they contain only a single Su(Hw) binding, both *eve* boundaries can promote transvection with the *gyspy* insulator as a pairing partner. In the case of *gypsy:homie*, we found that this interaction is orientation independent (Figure 11). These results could not be explained by a cohesin based loop extrusion mechanism. Nor would this mechanism provide an explanation for why *nhomie* and *homie* can partner in *trans* with the *gypsy* element but are unable to do so with a different CTCF dependent boundary *Fab-8*.

## Materials and methods

### Key resources table

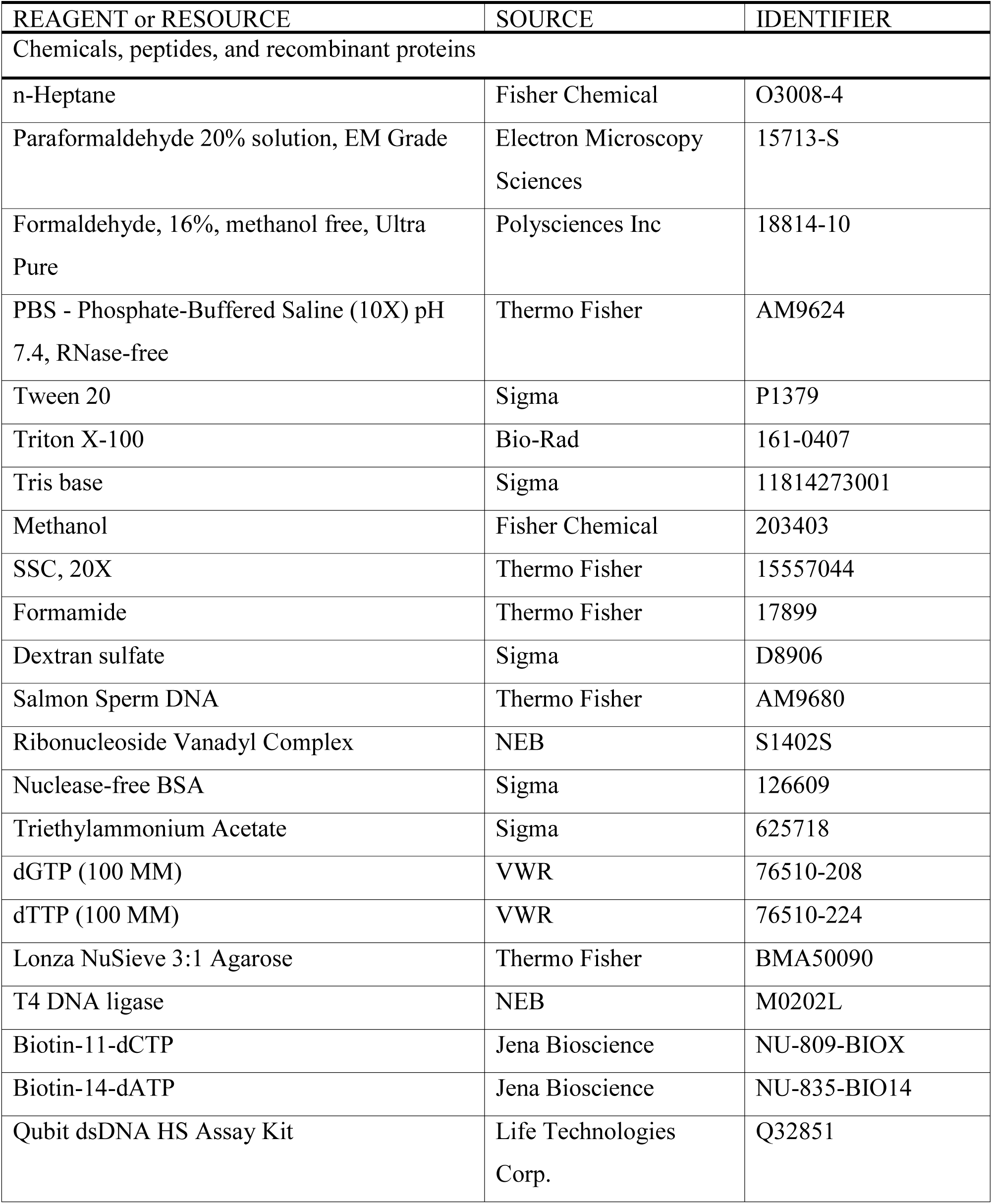
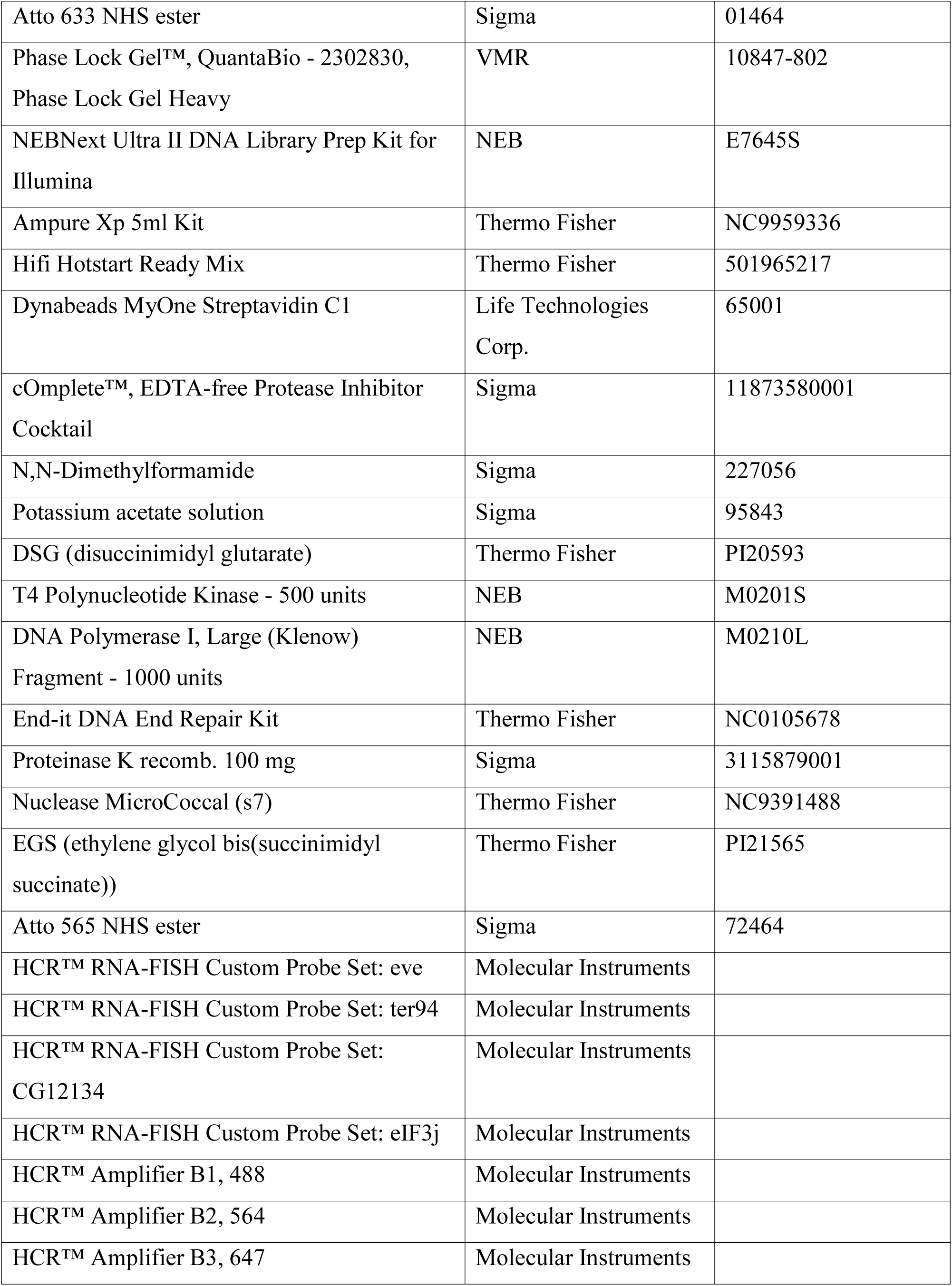
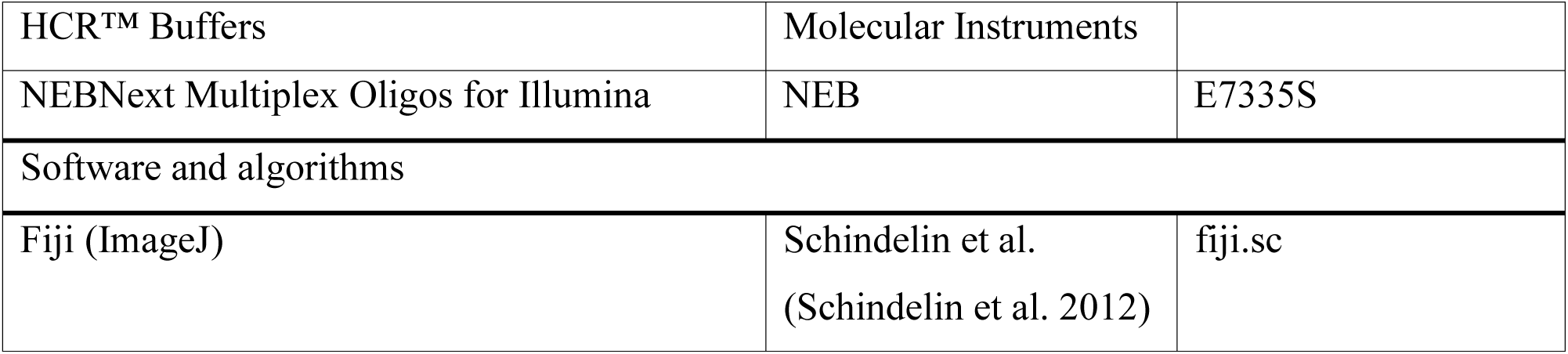

| REAGENT or RESOURCE | SOURCE | IDENTIFIER |
| --- | --- | --- |
| Chemicals, peptides, and recombinant proteins |  |  |
| n-Heptane | Fisher Chemical | O3008-4 |
| Paraformaldehyde 20% solution, EM Grade | Electron Microscopy Sciences | 15713-S |
| Formaldehyde, 16%, methanol free, Ultra Pure | Polysciences Inc | 18814-10 |
| PBS - Phosphate-Buffered Saline (10X) pH 7.4, RNase-free | Thermo Fisher | AM9624 |
| Tween 20 | Sigma | P1379 |
| Triton X-100 | Bio-Rad | 161-0407 |
| Tris base | Sigma | 11814273001 |
| Methanol | Fisher Chemical | 203403 |
| SSC, 20X | Thermo Fisher | 15557044 |
| Formamide | Thermo Fisher | 17899 |
| Dextran sulfate | Sigma | D8906 |
| Salmon Sperm DNA | Thermo Fisher | AM9680 |
| Ribonucleoside Vanadyl Complex | NEB | S1402S |
| Nuclease-free BSA | Sigma | 126609 |
| Triethylammonium Acetate | Sigma | 625718 |
| dGTP (100 mM) | VWR | 76510-208 |
| dTTP (100 mM) | VWR | 76510-224 |
| Lonza NuSieve 3:1 Agarose | Thermo Fisher | BMA50090 |
| T4 DNA ligase | NEB | M0202L |
| Biotin-11-dCTP | Jena Bioscience | NU-809-BIOX |
| Biotin-14-dATP | Jena Bioscience | NU-835-BIO14 |
| Qubit dsDNA HS Assay Kit | Life Technologies Corp. | Q32851 |
| Atto 633 NHS ester | Sigma | 01464 |
| Phase Lock Gel™, QuantaBio - 2302830,<br>Phase Lock Gel Heavy | VMR | 10847-802 |
| NEBNext Ultra II DNA Library Prep Kit for<br>Illumina | NEB | E7645S |
| Ampure Xp 5ml Kit | Thermo Fisher | NC9959336 |
| Hifi Hotstart Ready Mix | Thermo Fisher | 501965217 |
| Dynabeads MyOne Streptavidin C1 | Life Technologies<br>Corp. | 65001 |
| cOmplete™, EDTA-free Protease Inhibitor<br>Cocktail | Sigma | 11873580001 |
| N,N-Dimethylformamide | Sigma | 227056 |
| Potassium acetate solution | Sigma | 95843 |
| DSG (disuccinimidyl glutarate) | Thermo Fisher | PI20593 |
| T4 Polynucleotide Kinase - 500 units | NEB | M0201S |
| DNA Polymerase I, Large (Klenow)<br>Fragment - 1000 units | NEB | M0210L |
| End-it DNA End Repair Kit | Thermo Fisher | NC0105678 |
| Proteinase K recomb. 100 mg | Sigma | 3115879001 |
| Nuclease MicroCoccal (s7) | Thermo Fisher | NC9391488 |
| EGS (ethylene glycol bis(succinimidyl<br>succinate)) | Thermo Fisher | PI21565 |
| Atto 565 NHS ester | Sigma | 72464 |
| HCR™ RNA-FISH Custom Probe Set: eve | Molecular Instruments |  |
| HCR™ RNA-FISH Custom Probe Set: ter94 | Molecular Instruments |  |
| HCR™ RNA-FISH Custom Probe Set:<br>CG12134 | Molecular Instruments |  |
| HCR™ RNA-FISH Custom Probe Set: eIF3j | Molecular Instruments |  |
| HCR™ Amplifier B1, 488 | Molecular Instruments |  |
| HCR™ Amplifier B2, 564 | Molecular Instruments |  |
| HCR™ Amplifier B3, 647 | Molecular Instruments |  |
| HCR™ Buffers | Molecular Instruments |  |
| NEBNext Multiplex Oligos for Illumina | NEB | E7335S |
| Software and algorithms |  |  |
| Fiji (ImageJ) | Schindelin et al.<br>(Schindelin et al. 2012) | fiji.sc |

### Plasmid construction and transgenic lines

The dual reporters construct was described previously (Fujioka et al. 2016). In short, each reporter contains the *eve* basal promoter (-275 to +106 bp relative to *eve* start site), either the *LacZ* (*eve-LacZ*) or *EGFP* (*eve-GFP*) coding region, and the *eve* 3’ UTR (+1300 to +1525 bp). These two reporters are divergently transcribed. Test fragments were then inserted between the two reporters. The fragments used here are: a 500 bp fragment from *lambda* phage DNA, a 367 bp wild-type or Su(Hw)-mutant *homie* fragment (CDEF) (Fujioka et al., 2025), and a 602 bp wild-type or Su(Hw)-mutant *nhomie* fragment.

The -142 kb attP landing site was described previously (Fujioka et al. 2009). It contains two attP target sites for phiC31 recombinase-mediated cassette exchange (RMCE) (Bateman et al. 2006) and *mini-white* as a marker. RMCE can result in the insertion of the transgene in either orientation, and all four possible insertions of the transgenes can be recovered. For the experiments in this paper, we analyzed all four possible insertions for transgenes containing the *nhomie* fragment. For two of these, *G-eimohn-L* and *G-nhomie-L*, the *eve-LacZ* reporter is on the *eve* side of transgene *nhomie*, while *eve-GFP* is on the *hebe* enhancer side of the transgene *homie*. For the other two, *L-eimohn-G* and *L-nhomie-G*, the *eve-GFP* reporter is on the *eve* side of transgene *homie*, while *eve-LacZ* is on the *hebe* enhancer side of the transgene *nhomie*. For the *nhomie* Su(Hw) mutant *nhomieΔSH*, we analyzed the *G-nhomieΔSH-L* insert, where the *eve-LacZ* reporter is on the *eve* side of the transgene. For the two *homie* transgenes that were analyzed, *G-eimoh-L* and *G-eimohΔSH-L* the *eve-LacZ* reporter is also on the *eve* side of the transgene. RCMC events were identified by loss of *mini-white*, and the orientation of each insert was determined by PCR.

Two transgenes were used for the transvection assay (Fujioka et al., 2016). The reporter transgenes are the same ones used in -142 kb assay described above. For enhancer transgenes, modified boundaries (described in figures) were flanked by the anal plate ring (APR) - mesodermal enhancers and two neuronal enhancers. The transvection assay was carried out so that *LacZ* in the dual reporter transgene and the APR-mesodermal enhancers was placed on the same side of the boundary. With this arrangement, *LacZ* is activated by APR and mesodermal enhancers. As described in the text, modified *homie*, nhomie, *lambda* DNA, and *gypsy* were inserted into these transgenes. The *homie* DNAs were the 367 bp CDEF fragment, the 271 bp DEF fragment and the Su(Hw) mutant fragment DΔSuEF (Fujioka et al., 2025). The *lambda* DNA and *gypsy* fragments were also described previously (2025).

### HCR-FISH and digoxigenin *in situ* procedure

The sequences of target genes were obtained from Flybase (https://flybase.org)(Gramates et al. 2022). To design probes, the target gene sequences were submitted to the Molecular Instruments probe design platform (https://www.molecularinstruments.com/hcr-rnafish) (Choi et al. 2016), with parameters set to a 35 probe set size for *Drosophila melanogaster*. A similar method was designed based on published smFISH methods (Little and Gregor 2018; Trcek et al. 2017). 100-200 flies were placed in a cage with an apple juice plate at the bottom of the cage. For early stages, the embryos were collected for 7 hours, while for later-stage embryos, collections were overnight. Embryos from each plate were washed into collection mesh and dechorionated in bleach for 2min, then fixed in 5mL of 4% paraformaldehyde in 1X PBS and 5mL of heptane for 15min with horizontal shaking. The paraformaldehyde was then removed and replaced with 5mL methanol. The embryos were then devitellinized by vortexing for 30s, and washed in 1mL of methanol twice. Methanol was then removed and replaced by PTw (1X PBS with 0.1% Tween-20) through serial dilutions of 7:3, 1:1, and 3:7 methanol:PTw. The embryos were washed twice in 1mL of PTw and pre-hybridized in 200μL of probe hybridization buffer for 30min at 37°C. 0.4pmol of each probe set were added to the embryos in probe hybridization buffer, and the embryos were incubated at 37°C for 12-14h. The embryos were then washed 3X with probe wash buffer at 37°C for 30min and 2X with 5X SSCT(5X SSC+0.1% tween) at room temperature for 5min. Then the embryos were pre-amplified with 300μL amplification buffer for 10min at 25°C. Meanwhile, 6pmol of hairpin h1 and h2 were snap-cooled separately (95°C for 90s, cool to RT with a 0.1°C drop per second), and then mixed in 100μL of amplification buffer at room temperature. After that, the pre-amplification solution was removed from the embryos, and 100μL of hairpin h1/h2 mix were added to the embryos. Next, the embryos were incubated for 12-14h at room temperature in the dark. To remove excess hairpins, the embryos were washed in SSCT as follows: 2X for 5min, 2X for 30min, and 5X for 5min. Then, the embryos were washed with 1mL PTw for 2min and stained with DAPI/Hoechst at 1μg/mL for 15min at room temperature in the dark. The embryos were then washed with PTw 3X for 5min. Finally, the embryos were mounted on microscope slides with Vectashield and a #1.5 coverslip for imaging.

The procedure for digoxigenin *in situ* was described previously (Fujioka et al. 2025).

### Imaging, image analysis, and statistics

Embryos from HCR-FISH were imaged using a Nikon A1 confocal microscope system, with a Plan Apo 20X/0.75 DIC objective. Z-stack images were taken at interval of 2μm, 4X average, 1024x1024 resolution, and the appropriate laser power and gain were set for the 405, 488, 561, and 640 channels to avoid overexposure. Images were processed by ImageJ, and the maximum projection was applied to each of the stack images. To determine the presence of stripes in early embryos, multi-channel images were first split into single channels, and the stripe signal was then highlighted and detected by the MaxEntropy thresholding method. GraphPad Prism was used for data visualization and statistical analysis. Two-way ANOVA with Tukey’s multiple comparisons test for each pair of groups was used to determine the statistical significance for the percentage of embryos carrying stripes in *eIF3j* and *TER94* channels in each group.

### Imaging, image analysis and statistics

Embryos from smFISH were imaged using a Nikon A1 confocal microscope system with a Plan Apo 20X/0.75 DIC objective. Z-stack images were taken at intervals of 2μm, 4X average, 1024x1024 resolution, and the appropriate laser power and gain were set for the 405, 561, and 640 channels to avoid overexposure. Images were processed using ImageJ, and the maximum projection was applied to each of the stack images. To measure the stripe intensity of early embryos, multi-channel images were first split into single channels, and the stripe signal was then highlighted and selected by the MaxEntropy thresholding method. For APR intensity measurements, the ROI tool was used to crop out the APR region of late-stage embryos. The cells with APR signal were also highlighted and selected by MaxEntropy thresholding. The particle measurement tool was used to measure the average intensity of all cells that had a signal. At the same time, the background signal (average intensity) was taken from cells without a signal in the same embryo. The relative intensity (signal to background) for each embryo was calculated using the stripe signal and background from the same embryo. To make comparisons between independent biological replicates, the average background signal of all embryos from each replicate was calculated. The relative intensity of each embryo from each replicate was normalized based on the average background signal of all embryos from that replicate. GraphPad Prism was used for data visualization and statistical analysis. To compare intensity from embryos in different groups, different signals from the same embryo (e.g., *LacZ* and *GFP*) were paired, and paired two-tailed t-tests were used to calculate p-values. All raw measurements and normalized data are included in Supplemental Data S2.

### MicroC library construction

Embryos were collected on yeasted apple juice plates in population cages for 4 hours, incubated for 12hr at 25℃, then subjected to fixation as follows. Embryos were dechorionated for 2min in 3% sodium hypochlorite, rinsed with deionized water, and transferred to glass vials containing 5 mL PBST (0.1% Triton-X100 in PBS), 7.5 mL n-heptane, and 1.5mL fresh 16% formaldehyde. Crosslinking was carried out at room temperature for exactly 15min on an orbital shaker at 250rpm, followed by addition of 3.7 mL 2M Tris-HCl pH7.5 and shaking for 5min to quench the reaction. Embryos were washed twice with 15 mL PBST and subjected to secondary crosslinking. Secondary crosslinking was done in 10mL of freshly prepared 3mM final DSG and ESG in PBST for 45 min. at room temperature with passive mixing. The reaction was quenched by addition of 3.7mL 2M Tris-HCl pH7.5 for 5min, washed twice with PBST, snap-frozen, and stored at -80℃ until library construction.

Micro-C libraries were prepared as previously described (Batut et al. 2022) with the following modifications: 50uL of 12-16hr embryos were used for each biological replicate. 60U of MNase were used for each reaction to digest chromatin to a mononucleosome:dinucleosome ratio of 4. Libraries were barcoded, pooled, and subjected to paired-end sequencing on an Illumina Novaseq S1 100 nt Flowcell (read length 50 bases per mate, 6-base index read).

### Micro-C data processing

MicroC data for *D. melanogaster* were aligned to custom genomes edited from the Berkeley Drosophila Genome Project (BDGP) Release 6 reference assembly (dos Santos et al. 2015) with BWA-MEM (Li and Durbin 2009) using parameters **-S -P -5 -M**. Briefly, the custom genomes are simply insertions of the transgenic sequence into the –142kb integration site, as predicted from perfect integration. These events were confirmed using PCR post-integration. The resultant BAM files were parsed, sorted, de-duplicated, filtered, and split with Pairtools (https://github.com/mirnylab/pairtools). We removed pairs where only half of the pair could be mapped, or where the MAPQ score was less than three. The resultant files were indexed with Pairix (https://github.com/4dn-dcic/pairix). The files from replicates were merged with Pairtools before generating 100bp contact matrices using Cooler (Abdennur and Mirny 2020). Finally, balancing and Mcool file generation was performed with Cooler’s Zoomify tool.

Virtual 4C profiles were extracted from individual replicates using FAN-C (Kruse et al. 2020) at 400bp resolution. The values were summed across replicates and smoothed across three bins (1.2kb). Viewpoints were determined based on the most informative region for interpretation. We used an 800bp window located downstream of the *eve* promoter, in the gene body of either *GFP* or *LacZ*. For the *nhomie* and *homie* viewpoints, the 800 bp window centered on the boundary; however, due to the masking of the duplicated boundary and *eve* promoters in the data analysis, a reduced signal is expected (Fig. 9).

### Calculation and comparison of the insulation score at the *hebe* insertion site

We used the FAN-C (https://github.com/vaquerizaslab/fanc/tree/main) package to compute insulation scores on the merged Mcool maps corresponding to all the conditions, using a window size of 4 kb. The insulation score is defined as the log (base 2)-fold change between the signal within a 4 kb window centered on a site over the mean of all signal values along chromosome 2R. First, we computed boundaries using the boundary calling tool from FAN-C and identified the *hebe* insertion site for all mutants. Using these centered coordinates, we computed the insulation score at the *hebe* insertion site from the balanced contact maps. We then normalized this score relative to the local region around the *hebe* insertion site by subtracting the original insulation score from the mean of the insulation scores in a 40.2 kb window centered around the *hebe* insertion site. This mean-scaled normalization can be interpreted as the level of insulation at the *hebe* insertion relative to its local neighborhood. We computed the change in the normalized insulation between *G-lambda-L* and all mutants.

### Quantification of total contacts between the *hebe* transgene reporter and the *eve* locus

For all the mutants, we identified the location of the *hebe* locus and the *eve* locus (specifically, the region between *nhomie* and *homie*). We then visualized the interaction region between the *hebe* insertion and the *eve* locus to manually segment bounding boxes of interaction between the *LacZ* reporter and the *eve* locus. After matrix balancing and masking of null values, we took the mean of the contact frequencies within this bounding box for each dataset. This became the average contact frequency between *LacZ* and *hebe* transgene. We repeated this procedure for *gfp*.

## Supporting information

All supplemental figures

## Data availability

Sequence data are available at GEO (accession number GSE330020). Analysis and notes are available at https://github.com/pritykinlab/suhw-tad-boundary-mechanisms.

## Acknowledgements

Part of this work was supported by NIH grants to Paul Schedl (5R35GM126975), to James B. Jaynes (1R01GM137062) and to Yuri Pritykin (DP2AI171161). Princeton Bioengineering Biocondensates Program to Yuri, and a New Jersey Commission on Cancer Research (COCR23PDF011) to Wenfan Ke. Authors would like to thank Gordon Grey for preparing fly food, members of the Lewis Sigler Genomics Core facility for the assistance with DNA sequencing and Qing Liu for excellent technical assistance.

