## Supplementary material for "Shared binding sites for the chromosomal architectural protein Su(Hw) mediate physical interactions between *Drosophila* TAD boundaries": All supplemental figures

**Figure supplemental**

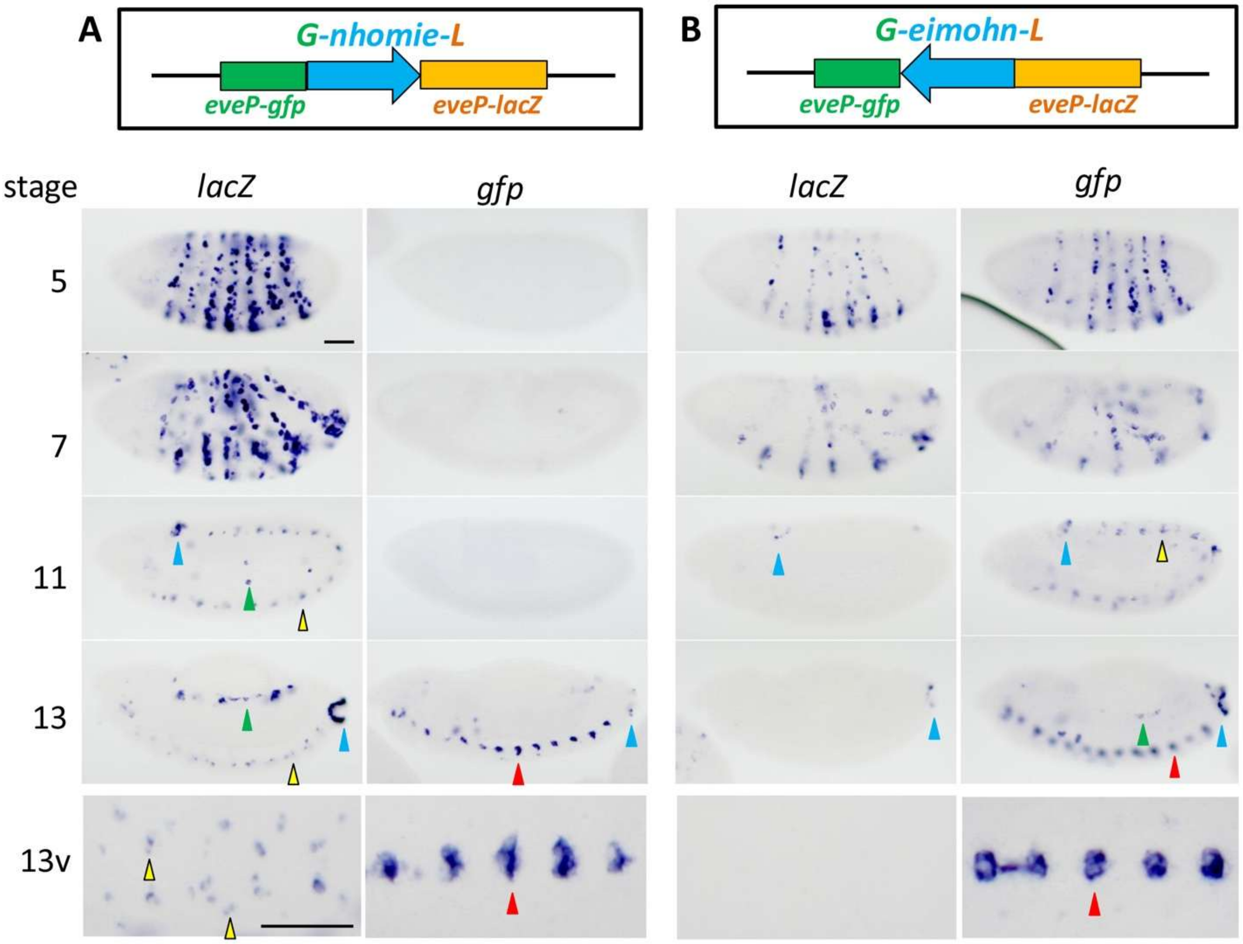


**Figure 1-figure supplemental 1 Digoxigenin *in situ* hybridization showing expression of *lacZ* and GFP mRNAs from the *G-nhomie-L* (A) and *G-eimohn-L* (B) transgenes.** The transgene constructs are illustrated on the top. Embryonic stages 5, 7, 11, 13 are shown. Ventral views of stage 13 (13v) are shown in the bottom row. Arrowheads indicate followings; Blue: anal plate ring (APR), Yellow: *eve* positive neurons, Green: mesoderm, and Red: *hebe* midline cell cluster. Scale Bar: 50μm.

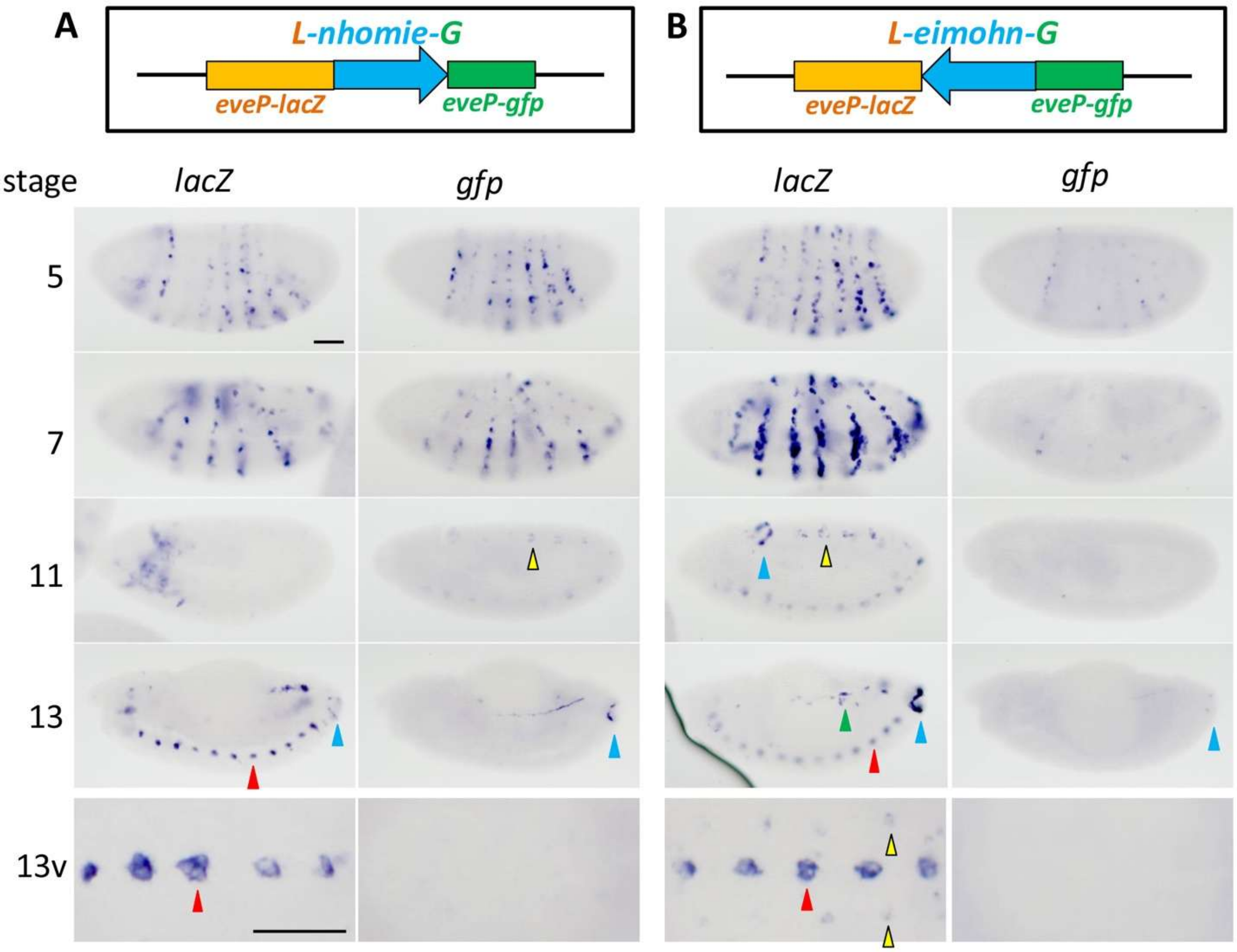


**Figure 1-figure supplemental 2 Digoxigenin *in situ* hybridization showing expression of *lacZ* and GFP mRNAs from the *L-nhomie-G* (A) and *L-eimohn-G* (B) transgenes.** The transgene constructs are illustrated on the top. Embryonic stages 5, 7, 11, 13 are shown. Ventral views of stage 13 (13v) are shown in the bottom row. Arrowheads indicate followings; Blue: anal plate ring (APR), Yellow: *eve* positive neurons, Green: mesoderm, and Red: *hebe* midline cell cluster. Scale Bar: 50µm.

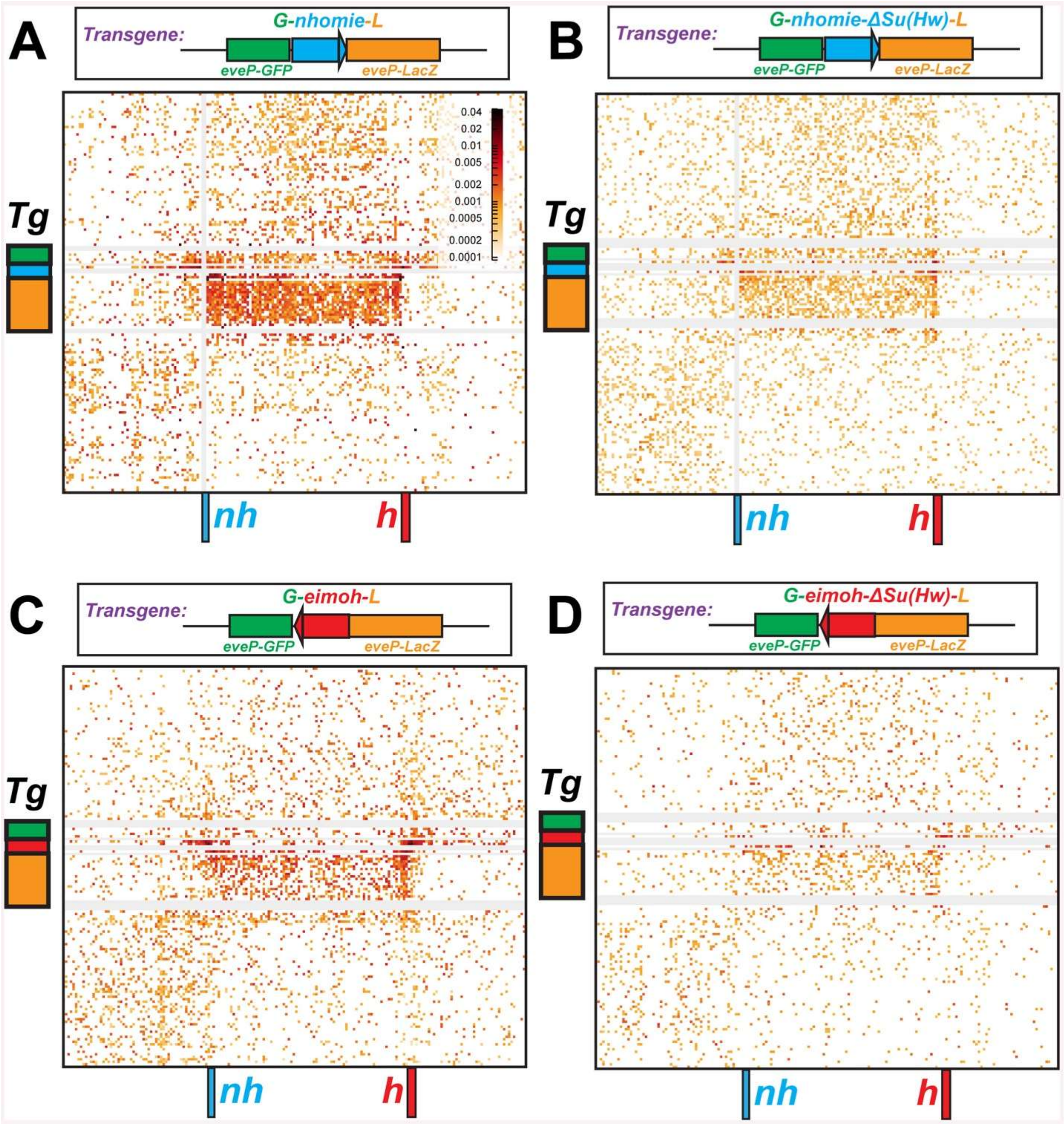


**Figure 7-figure supplemental 1. Blow up of the MicroC interaction profiles between transgenes inserted at -142 kb and the *eve* TAD and TADs flanking the *eve* TAD**. A) *G-nhomie-L* B) *G-nhomieΔSu(Hw)-L* C) *G-eimoh-L* and D) *G-eimohΔSu(Hw)-L*. Position of transgene indicated on right, while the positions of the *nhomie* (*nh*) and *homie* (*h*) boundaries indicated below.

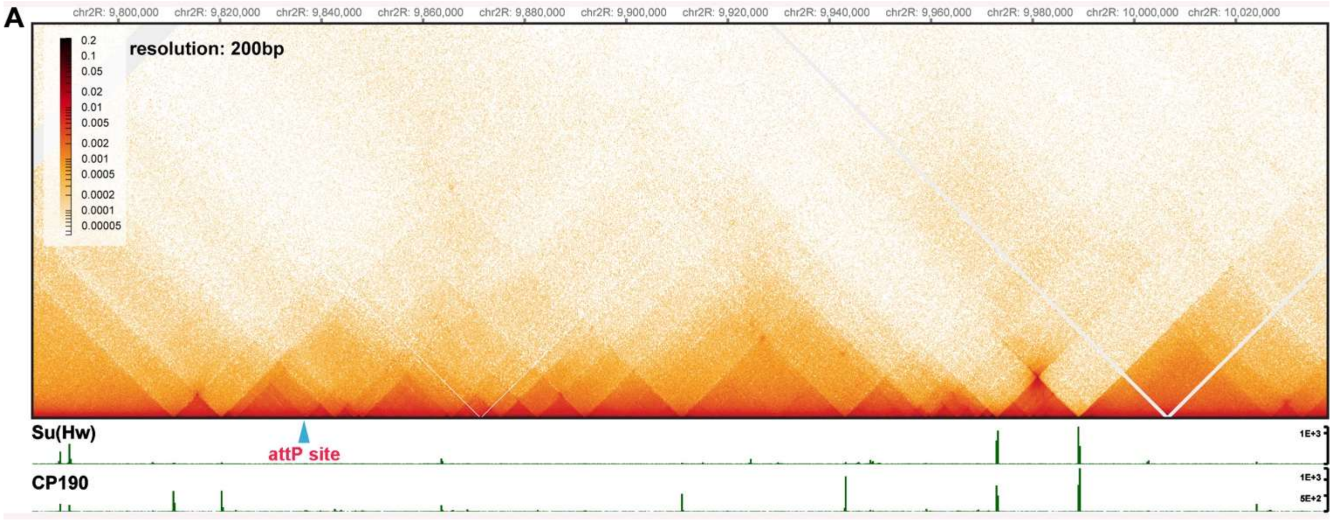


**Figure 7-figure supplemental 2. Su(Hw) protein and TADs near the *hebe* attP site and the *eve* TAD** Location of TADs and Su(Hw) protein in region containing the *hebe* attP site and the *eve* TAD.

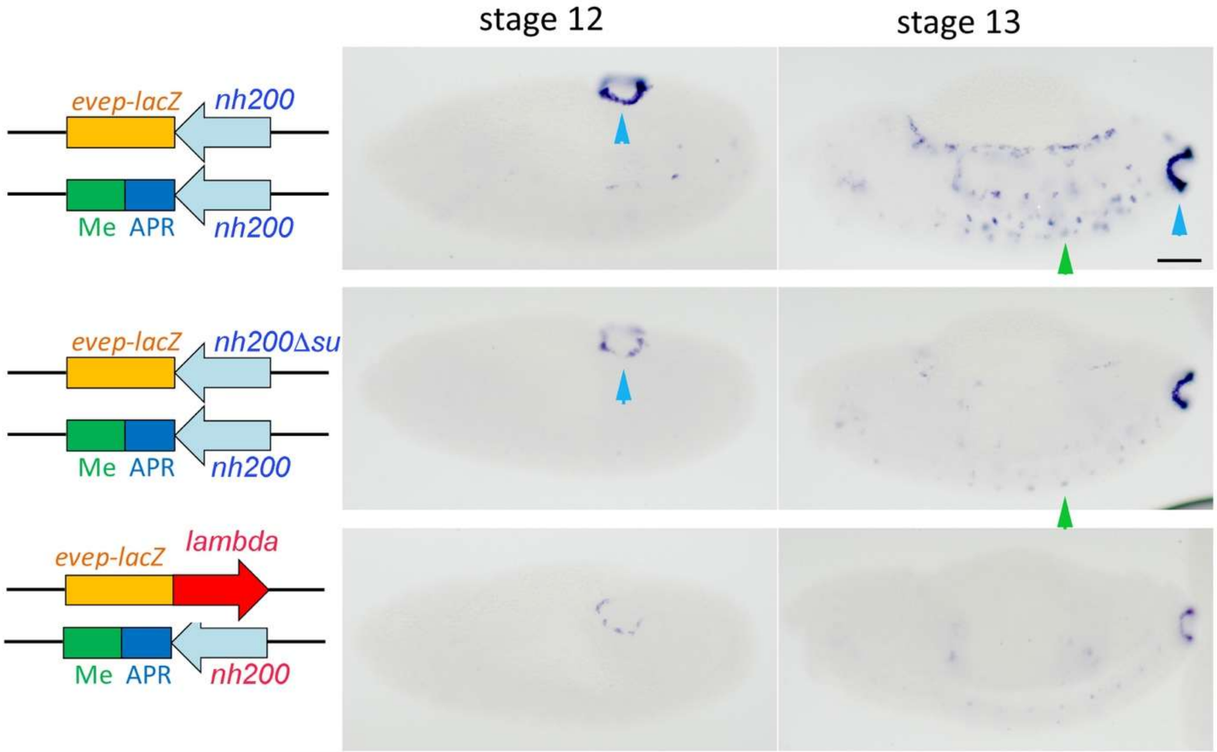


**Figure 10-figure supplemental 1. *nhomie*-dependent transvection is weakened but not eliminated by loss of the Su(Hw) binding site.** Digoxigenin *in situ* hybridization showing *LacZ* expression in the transvection assay. The transgene constructs used to assay transvection are shown in the diagrams on the left. The negative control has a 500 bp *lambda* DNA fragment. In this experiment we used a smaller *nhomie* fragment of 200 bp, instead of the larger 600 bp fragment that was used in the -142 kb assay and in the Figure 10 figure supplemental 1. The long-distance activity of this smaller *nhomie* fragment is greatly reduced compared to larger 600 bp *nhomie* fragment. Stages 12, and 13 are shown. Arrowheads indicate the following: blue: APR and green: mesoderm. Blue: anal plate ring (APR), and Green: mesoderm. Scale bar: 50µm.

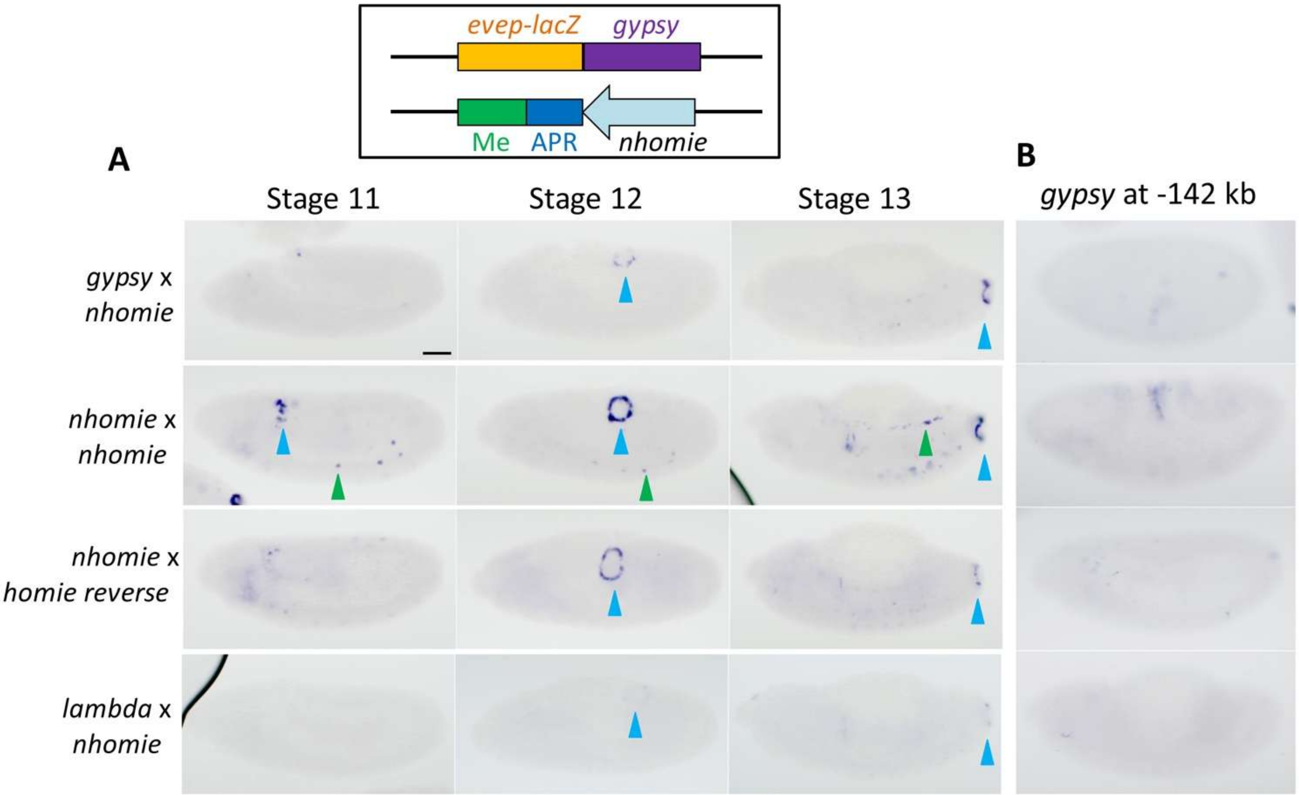


**Figure 11-figure supplemental 1. The su(Hw) *gypsy* insulator supports transvection with *nhomie*.** A) Digoxigenin *in situs* showing expression of *lacZ* in the transvection assay. The *nhomie* and *gysyp* transgene constructs are illustrated on the top. The transgenes used are indicated as reporter x enhancer at the left. A 500 bp *lambda* DNA fragment was used as negative control (*lambda*). The 349 bp *gypsy* su(Hw) insulator fragment and 602 bp *nhomie*, as well as *homieFEDC* were tested. Stages 11, 12 and 13 are shown. Arrowheads indicate followings; Blue: anal plate ring (APR), and Green: mesoderm. Scale Bar: 50µm. B: Digoxigenin *in situ* hybridization of *lacZ* expression when the dual reporter inserted in the -142 kb attP site contains the 349 bp *gypsy* insulator fragment.
